# Drug inhibition and substrate alternating flipping mechanisms of human VMAT2

**DOI:** 10.1101/2024.02.28.582500

**Authors:** Feiwen Wei, Huihui Liu, Wei Zhang, Jufang Wang, Yanqing Zhang

**Affiliations:** Shanghai Fifth People’s Hospital, Fudan University, and Shanghai Key Laboratory of Medical Epigenetics, International Co-laboratory of Medical Epigenetics and Metabolism (Ministry of Science and Technology), Institutes of Biomedical Sciences, Fudan University, Shanghai 200032, China; Warshel Institute for Computational Biology, School of Medicine, The Chinese University of Hong Kong, Shenzhen, Guangdong 518172, China

**Author notes:** These authors contributed equally (Feiwen Wei, Huihui Liu, Wei Zhang). Correspondence (Y.Z.).

## Abstract

Vesicular monoamine transporters (VMAT1/2) are responsible for loading and packaging monoamine neurotransmitters into synaptic vesicles, including serotonin (5-HT), dopamine (DA), norepinephrine, and histamine. Dysregulation of VMAT2 within the central nervous system can lead to schizophrenia, mood disorders, and Parkinson’s disease, due to the imbalances of these monoamine neurotransmitters. Medications such as tetrabenazine (TBZ) and valbenazine (VBZ) targetting VMAT2 are approved for treating chorea associated with Huntington’s disease and Tardive Dyskinesia. Our cryo-EM studies and molecular dynamics (MD) simulations on VMAT2 bound to drug inhibitors (TBZ and VBZ) and substrates (5-HT and DA), unveil the inhibition mechanism of VMAT2, alternating flipping mechanism of substrates during loading, translocation, and release, as well as the interplay between protonation of crucial acidic residues and substrate release. These findings enhance the understanding of VMAT-mediated monoamine neurotransmitter transport, fostering drug development for neurological and neuropsychiatric disorders, with a specific emphasis on VMATs.

## Introduction

Vesicular amine transporters, also known as solute carrier 18 (SLC18) family members, are responsible for loading and packaging amine neurotransmitters into synaptic vesicles^1-3^. The SLC18 family comprises vesicular monoamine transporters 1/2 (VMAT1/2, encoded by *SLC18A1/2*), vesicular acetylcholine transporter (VAChT, encoded by *SLC18A3*), and vesicular polyamine transporter (VPAT, encoded by *SLC18B1*) (Fig. S1C)^4-7^. These transporters play indispensable roles in the precise regulation of packaging, storage, and controlled release of neurotransmitters^8,9^. The vesicular monoamine transporters, VMAT1/2, are responsible for the uptake of monoamine neurotransmitters, including serotonin (5-HT), dopamine (DA), norepinephrine, and histamine, into synaptic vesicles^3^. VMAT1 is primarily distributed in peripheral tissues, especially in neuroendocrine cells, while VMAT2 is predominantly expressed in neurons within the central nervous system^10-13^.

These monoamine neurotransmitters, such as 5-HT and DA, are critical signal molecules within the central nervous system, participating in a broad range of physiological processes, encompassing emotion, cognition, motor, and learning^14-17^. Dysfunctional VMAT2 can lead to a spectrum of neurological and neuropsychiatric disorders, such as schizophrenia, mood disorders, and Parkinson’s disease^18-23^. Consequently, VMAT2 has emerged as a target of many drugs for neurological and neuropsychiatric disorders, such as reserpine, tetrabenazine (TBZ) and its derivatives^24-26^. TBZ was first synthesized in 1950 to treat psychoses and schizophrenia, and it was the first FDA-approved medication for chorea associated with Huntington’s disease in 2008^27^. Some derivatives of TBZ have also been produced as VMAT inhibitors, such as deutetrabenazine (dTBZ) and valbenazine (VBZ)^28^. VBZ has received FDA approval for the treatment of Tardive Dyskinesia in adults^29,30^.

VMAT2 operates as a secondary active transporter, relying on the cytoplasm-directed proton electrochemical gradient established by the proton pump vesicular H^+^-ATPase^31,32^. To unravel the structural intricacies of substrate recognition and drug inhibition, we determined the cryo-EM structures of VMAT2 bound to substrates, 5-HT and DA, and inhibitors, TBZ and VBZ. In combination with MD simulations and functional studies of mutagenesis, our work reveals the substrate transport mechanism and inhibition mechanism of human VMAT2.

## Results

### Structure determination

To surmount challenges encountered in determining the cryo-EM structures of the small membrane protein VMAT2, we implemented a methodology previously outlined in the literature^33^. This method involved the fusion of a BRIL-tag at the N-terminus of the VMAT2 (referred to as VMAT2_EM_), followed by incubation of the resultant fused protein with anti-BRIL Fab and anti-Fab nanobody to enlarge the particle diameter (Fig. S1A, B). We evaluated the transport activity of VMAT2_EM_, which preserved around 60% of the uptake activity observed in the wild-type VMAT2 (VMAT2_WT_). When exposed to TBZ and VBZ, VMAT2_EM_ displayed fully inhibited activity as well as VMAT2_WT_ (Fig. 1A). Therefore, VMAT2_EM_ recapitulates the transport and drug inhibition characteristics of VMAT2_WT_. We successfully solved the structures of VMAT2 bound to TBZ, VBZ, 5-HT, and DA at resolutions of 3.5 Å, 3.5 Å, 3.0 Å, and 3.6 Å, respectively (Fig. 1B-D; Figs. S2-5 and Table S1), without clear observations of BRIL, Fab and nanobody densities. The cryo-EM densities corresponding to TBZ, VBZ, and 5-HT exhibited high quality, allowing unambiguous assignment of these ligand poses, which were further validated by molecular dynamics (MD) simulations. While determining the precise model of DA from its density with lower resolution was challenging, we estimated the pose of DA with further validation through ligand-free docking and MD simulations (Fig. 1C, F; Figs. S6 and Table S2).

**Fig. 1.**
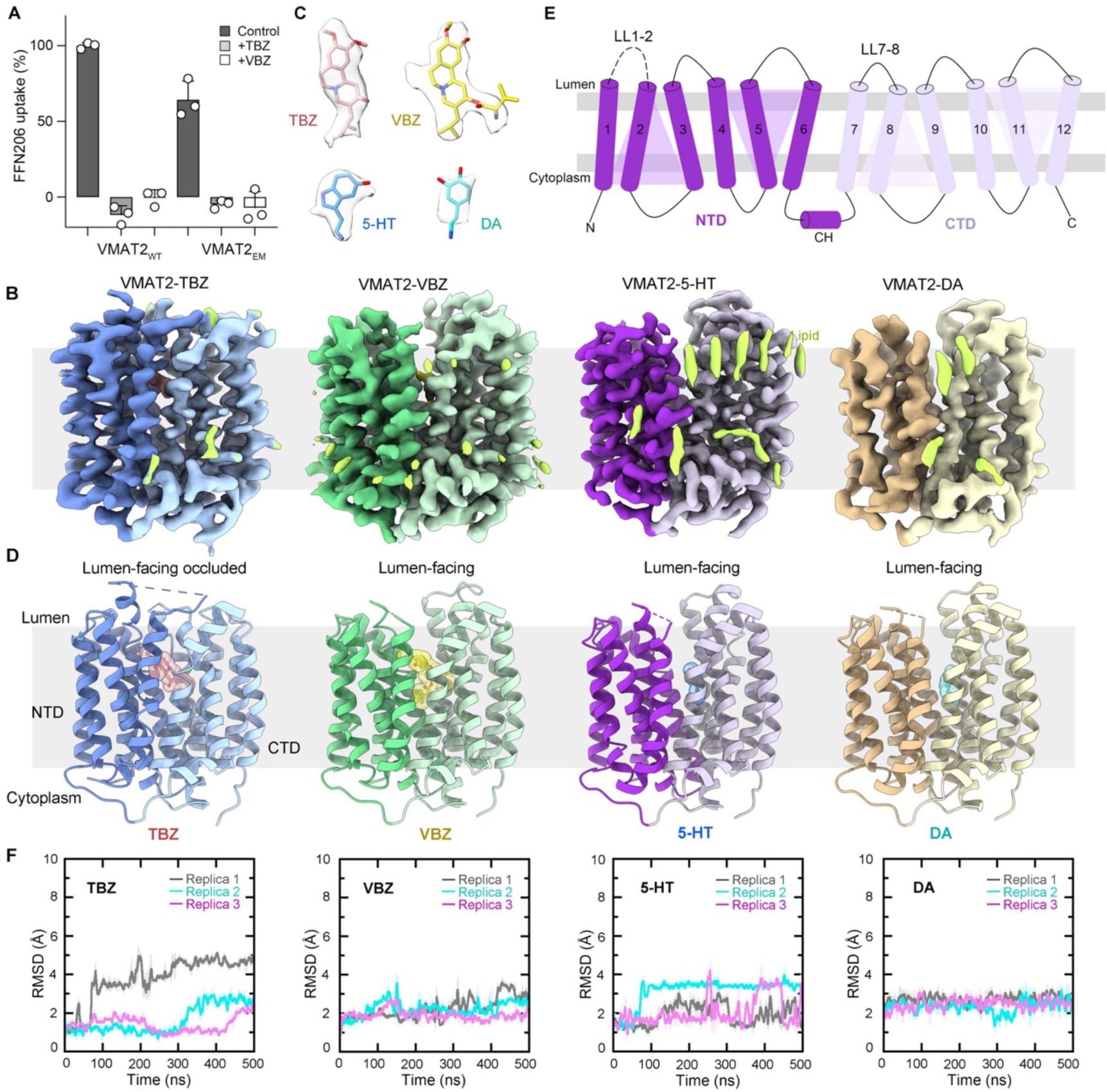
Structure determination and overall structures of human VMAT2 bound to substrates and inhibitors. **A** Fluorescent uptake of FFN206 and inhibition for wild-type transporter (VMAT2_WT_) and the cryo-EM construct (VMAT2_EM_) (mean ± SEM, *n* = 3 biologically experiments). **B** Cryo-EM density maps of VMAT2 bound to TBZ, VBZ, 5-HT, and DA. TBZ, tetrabenazine; VBZ, valbenazine; 5-HT, serotonin; DA, dopamine. **C** Cryo-EM densities of TBZ, VBZ, 5-HT, and DA. Density contour levels are 0.35, 0.23, 0.36, and 0.06 in ChimeraX, respectively. **D** Cartoon representations of structures of VMAT2 bound to TBZ, VBZ, 5-HT, and DA. **E** The topology of VMAT2. The transmembrane helices (TM) numbers are labeled, and the flexible region in our structure is shown as dashed line. The N-terminal domain (NTD), cytoplasmic helix (CH) are colored in deep purple, and the C-terminal domain (CTD) is colored in light purple. **F** Root mean square deviation (RMSD) plots of MD simulations for TBZ, VBZ, 5-HT, and DA in 500 ns for three replicas.

The structures show that VMAT2 is a compact membrane protein adopting the major facilitator superfamily (MFS) fold^34,35^ which comprises 12 transmembrane helices (TMs) organized into two structurally pseudo-symmetry domains: the N-terminal domain (NTD, TMs 1-6) and the C-terminal domain (CTD, TMs 7-12), with both termini situated in the cytoplasm. In our structural analysis, the extensive luminal loop between TM 1 and 2 (LL1-2) demonstrated flexibility and remained undetectable (Fig. 1E). Overall, VBZ, 5-HT, and DA bind to VMAT2 in the lumen-facing state. Notably, TBZ was found to ensnare VMAT2 in the lumen-facing occluded state (Fig. 1D).

### TBZ binding sites

TBZ captured VMAT2 in a lumen-facing occluded state that is inaccessible from both sides of the membrane (Fig. 2). The cytoplasmic gate is effectively sealed by two layers of residues: one is composed of M204, M221, M403, and Y422, and the other involves two salt bridges, D214-R357 and R217-D411, forming an interaction network with S416 and Y418 (Fig. 2B). Meanwhile, in conjunction with a series of hydrophobic residues—V41, P45, F135, and F334—W318 predominantly obstructs the luminal gate by acting as a plug, effectively blocking the luminal mouth (Fig. 2A, C, E, F). The induced intermediate occluded conformation of VMAT2 might be a key aspect of the inhibition mechanism of TBZ, which would dampen the transport cycle^36-38^. TBZ occupies the central cavity formed by TMs 1, 4, 7, and 10. The binding sites of TBZ predominantly consist of non-polar residues from both NTD and CTD, including L37, I44, L228, V232, and I308, compensating the hydrophobic skeleton of TBZ. The highly conserved residue E312 forms a salt bridge with the positively-charged tertiary amine of TBZ, situated approximately in the middle of TBZ, consistent with previous study that the equivalent residue (E313) in rat VMAT2 is crucial for inhibitor recognition and substrate uptake^39^. F135 engages in pi-pi stacking with the benzene ring of TBZ, and R189 establishes a polar interaction with one methoxy group of TBZ, contributing to extra stability to the binding of TBZ (Fig. 2C; Fig. S8A). It is noteworthy that TBZ exhibits selective binding to and inhibition of VMAT2 rather than VMAT1^40^, which might be attributed to residue differences in the TBZ binding sites, including L37, V232, I308, and Y433 in VMAT2 (Fig. S1C).

**Fig. 2.**
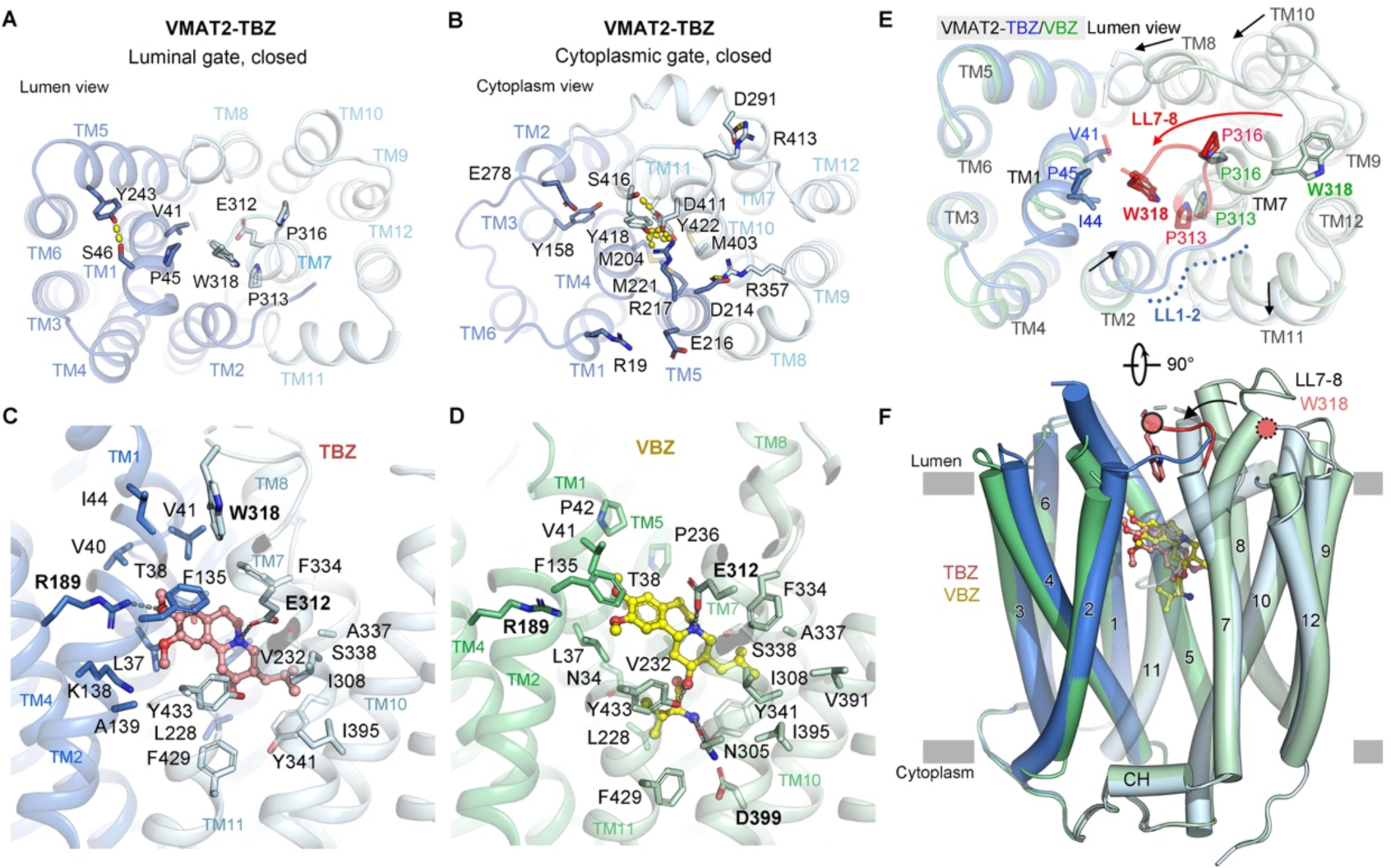
Inhibition mechanisms of TBZ and VBZ. **A** The luminal gate in the VMAT2-TBZ from the lumen view. **B** The cytoplasmic gate in the VMAT2-TBZ from the cytoplasm view. **C-D** Inhibitor binding sites of TBZ (C) and VBZ (D), in the lumen-facing occluded state and the lumen-facing state of VMAT2, respectively. **E** Conformational changes between TBZ- and VBZ-bound conformations of VMAT2 from the lumen view. **F** Superposition of VMAT2-TBZ (lumen-facing occluded, blue) and VMAT2-VBZ (lumen-facing open, green). W318 is highlighted in red.

### VBZ binding sites

As a derivative of TBZ, VBZ is the valine ester of (+)-α-dihydrotetrabenazine (HTBZ), which is the most active metabolite of TBZ^41^. In contrast to TBZ, VBZ trapped VMAT2 in a vesicular lumen-facing state. The cytoplasmic gate aligns with that of VMAT2-TBZ, while the luminal gate is open (Fig. 2F and Fig. S7A). In general, VBZ binds to the central cavity in a manner akin to TBZ, involving many overlapped non-polar residues that contribute to hydrophobic interactions with VBZ. E312 is also found to engage in a salt bridge with the positively-charged tertiary amine of VBZ. However, slightly different from TBZ, structural superposition reveals that the position of VBZ undergoes a slight shift towards the lumen, possibly due to steric effect of the additional valine moiety in VBZ (Fig. 2F). This minor positional adjustment of VBZ, compared to TBZ, disrupts both the polar interaction between R189 and the methoxy group and the pi-pi stacking involving F135. Simultaneously, the amine group from the valine moiety of VBZ establishes polar interactions with N305 and Y341 proximal to D399 (Fig. 2D; Fig. S8B). These alterations in the binding sites between VBZ and TBZ might be potential factors contributing to the distinctive conformations observed in VMAT2.

### Locking VMAT2 in different conformations by TBZ and VBZ

Upon structural superposition of VMAT2-TBZ/VBZ, a contracted movement is evident in TMs 2, 8, and 10 in VMAT2-TBZ. Specifically, the C-terminal of TM 7 undergoes a transition from a helix to a loop (luminal loop 7-8, LL7-8), leading to the repositioning of W318 to block the luminal opening. This transition coincides with a dilated motion of TM 11 and the emergence of the luminal loop 1-2 (LL1-2), which is inserted between LL7-8 and TM 11 (Fig. 2E). The entrapment of VMAT2 in different conformations by VBZ and TBZ suggests their involvement in different inhibition mechanisms. We propose that both VBZ and TBZ bind to VMAT2 in the lumen-facing conformation, but VBZ traps the transporter in the lumen-facing state, while TBZ induces conformational changes leading to the lumen-facing occluded state, thereby obstructing the transport cycle at different steps.

### Substrate binding sites of 5-HT

To explore the mechanism of monoamine neurotransmitter transport by VMAT2, we analyzed the cryo-EM structure bound to 5-HT. 5-HT captured VMAT2 in the vesicular lumen-facing conformation (Fig. 1D). The cytoplasmic gate is sealed in a manner similar to VBZ-bound conformation (Fig. S7). Even though the cryo-EM density of 5-HT supports the pose where its amine group points towards D399, referred to as Pose_N-D399_, the ligand-free docking suggested the existence of an alternate flipped pose where the amine group points to E312, named Pose_N-E312_. To confirm the pose of 5-HT, we conducted MD simulations, revealing that 5-HT in Pose_N-D399_ was stable, while 5-HT in Pose_N-E312_ rapidly flipped to Pose_N-D399_ within 4 ns in three replicas (Fig. S6). Based on the cryo-EM density of 5-HT and the results of MD simulations, we confidently constructed 5-HT with the pose of Pose_N-D399_ in the lumen-facing state. The positively-charged amine group of 5-HT establishes a salt bridge with D399 and forms polar interactions with N305, Y341, and Y433. The hydroxyl group of 5-HT associates with E312 through hydrogen-bond interactions. Additionally, V232, I308, and F334 contribute to hydrophobic interaction with the indole group of 5-HT (Fig. 3A; Fig. S8C).

**Fig. 3.**
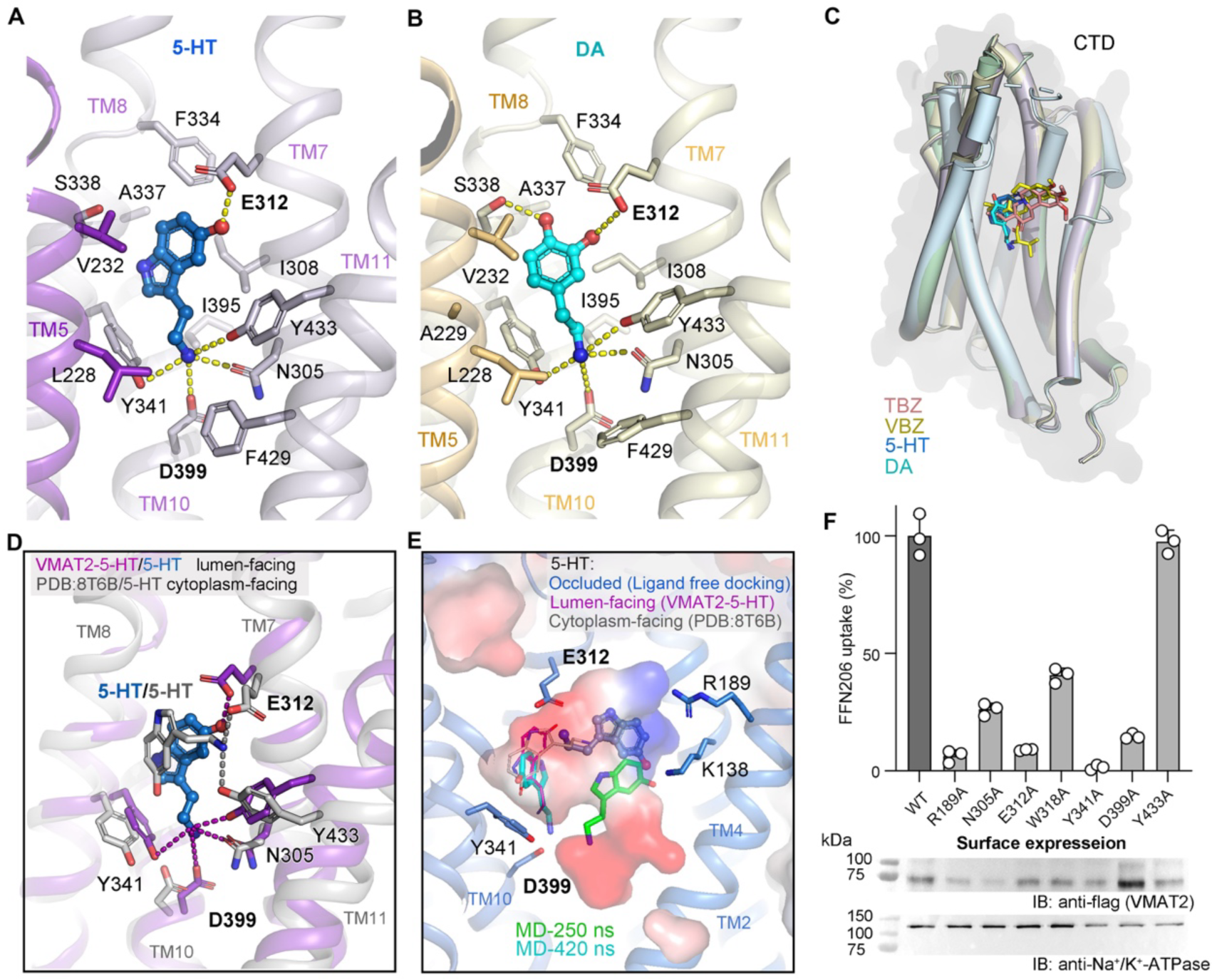
Substrate binding mechanisms of 5-HT and DA. **A-B** Substrate binding sites of 5-HT (A) and DA (B) in the lumen-facing state of VMAT2. **C** Superposition of all four ligand-binding sites in VMAT2 in this study. Only the C-domains are shown. TBZ, VBZ, 5-HT and DA are in salmon, yellow, marine, and cyan, respectively. **D** Superposition of the lumen-facing (5-HT-bound, this study, purple and blue) and cytoplasm-facing (5-HT-bound, PDB: 8T6B, gray) conformations of VMAT2. **E** Representative configurations of 5-HT in the lumen-facing occluded state of VMAT2, including one pose predicted with Maestro (marine), and two poses obtained from MD simulations (green and cyan, respectively). The lumen-facing occluded protein (marine) is taken from VMAT2-TBZ. The structure of 5-HT in lumen-facing state (purple) and cytoplasm-facing state (PDB: 8T6B, gray) are shown as references. **F** Fluorescent uptake of FFN206 for wild-type VMAT2 and mutants (mean ± SEM, *n* = 3 biologically experiments) (upper panel). Measurements of cell surface expression levels of VMAT2_WT_ and mutants by immunoblot with Na^+^/K^+^-ATPase as the internal control (lower panel).

### Substrate binding sites of DA

The binding of DA traps VMAT2 in the lumen-facing state similar to 5-HT-bound state (Fig. 1D; Fig. S7B). However, we encountered challenges in determining the pose of DA due to its lower resolution. The ligand-free docking results of Maestro revealed a ratio of approximately 2:1 for Pose_N-D399_: Pose_N-E312_ for DA. In consistence with 5-HT, MD simulations indicated that DA in Pose_N-D399_ was stable and DA in Pose_N-E312_ rapidly flipped to Pose_N-D399_ within 29 ns in three replicas, supporting the Pose_N-D399_ for DA (Fig. S6). Consequently, we built DA in Pose_N-D399_ in the lumen-facing state of VMAT2. The binding sites of DA closely resemble those of 5-HT. Additionally, the two hydroxyl groups are associated with E312 and S338, respectively (Fig. 3B; Fig. S8D). The cell-based fluorescent FFN206 uptake assay ^42^ demonstrated that mutations of N305A, E312A, Y341A, and D399A resulted in a significant decrease in transport activity, underscoring their essential roles in transport (Fig. 3F), in accord with previous studies suggested these residues are crucial for substrate transport and drug recognition^39,43,44^. Considering that the shared monoamine features of the substrates of vesicular monoamine transporters and the common binding mode of 5-HT and DA, the highly conserved E312 and D399 likely play key roles in the transport of monoamine neurotransmitters (Fig. S1).

### Drug inhibition mechanism

The binding sites of TBZ/VBZ overlap substantially with those of 5-HT/DA, and structural superposition reveals an occupancy overlap between the isobutyl group of TBZ/VBZ and the indole group of 5-HT or the benzene ring of DA (Fig. 3C; Fig. 5B). Intriguingly, the position of the amine group in the valine moiety of VBZ closely aligns with that of the amine group of 5-HT/DA, corresponding to the observation that almost all the binding sites of 5-HT/DA appear in the binding sites of VBZ. This suggests that VBZ might function as an inhibitor by acting as a substrate mimic, consistent with the observation that VBZ traps VMAT2 in the lumen-facing state, similar to 5-HT/DA, while TBZ induces an intermediate state. Considering the molecular difference between VBZ and TBZ, the binding of monoamine substrate in Pose_N-D399_ might trigger conformational changes of VMAT2 from the lumen-facing occluded state to the lumen-facing state during transport.

**Fig. 4.**
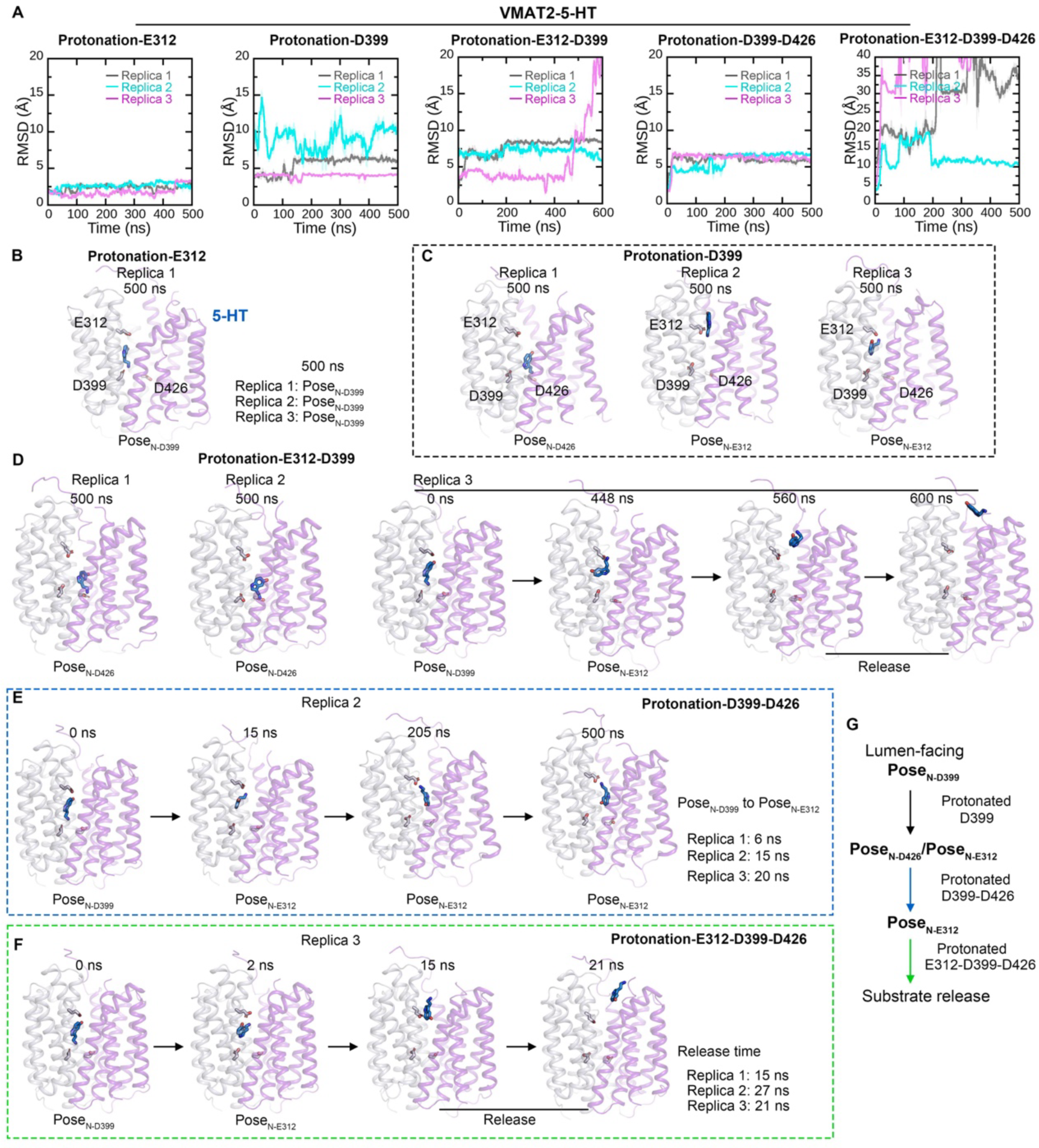
Molecular dynamics (MD) simulations of VMAT2-5-HT structure in different cases of protonation. **A** Root mean square deviation (RMSD) plots of MD simulations for the 5-HT-bound lumen-facing VMAT2 structure, with only E312 protonated, only D399 protonated, both E312 and D399 protonated, both D399 and D426 protonated, or all E312, D399 and D426 protonated. **B-F** MD snapshots of the 5-HT-bound lumen-facing VMAT2 structure, with only E312 protonated (B), only D399 protonated (C), both E312 and D399 protonated (D), both D399 and D426 protonated (E), and all E312, D399 and D426 protonated (F). Frames are selected as shown. 5-HT, serotonin. **G** Schematic of the substrate pose change versus the protonation site during substrate release.

**Fig. 5.**
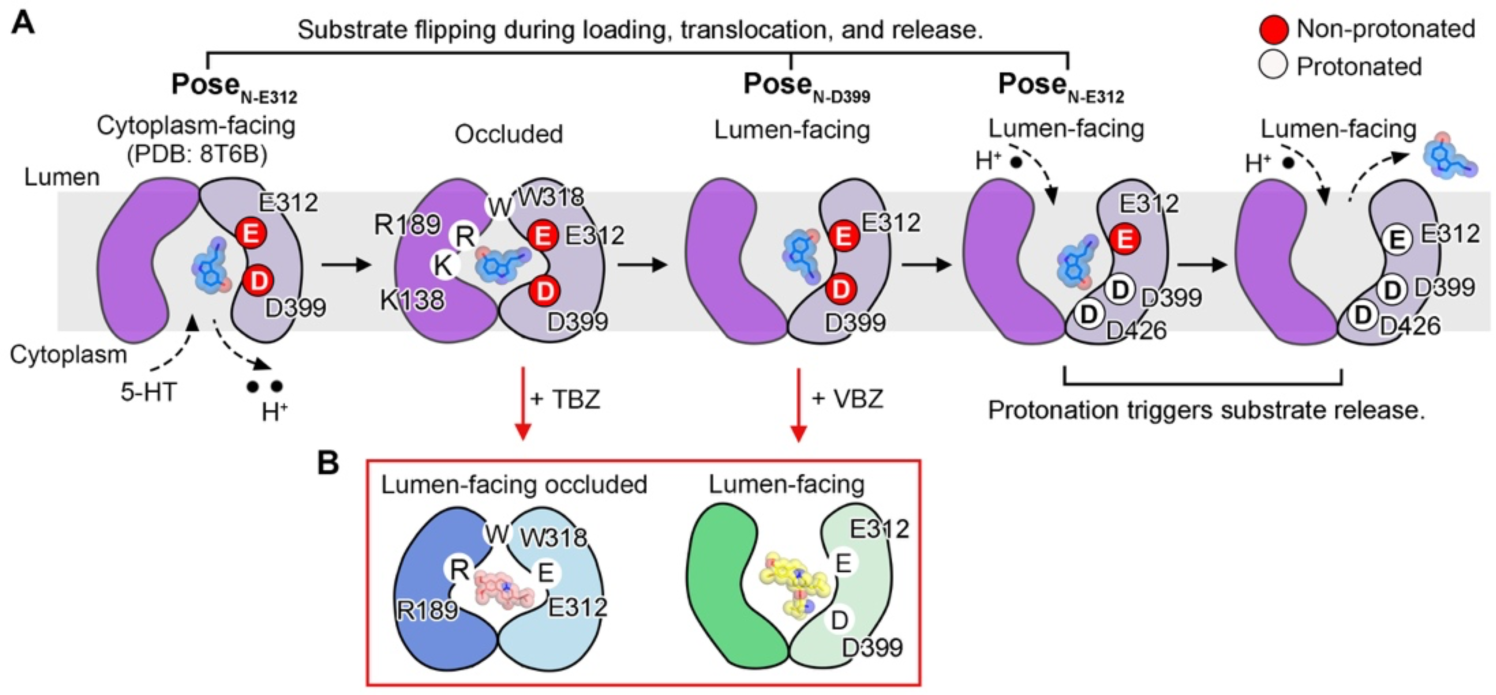
Schematic of substrate transport and drug inhibition mechanisms of VMAT2. **A** Schematic illustrating the substrate flipping mechanism during transport in VMAT2, using 5-HT as an example. Substrate experiences flipping alterations between Pose_N-E312_ and Pose_N-D399_ during substrate loading, conformational changes, and substrate release. Subsequent protonations of D399 and E312 initiate substrate release, a process significantly accelerated by D426 protonation as indicated by MD simulations. **B** The drug inhibitors, TBZ and VBZ, trap VMAT2 in the lumen-facing occluded state and lumen-facing state, respectively.

### Substrate flipping mechanism during transport

In line with recently reported structures of VMAT2-5-HT^45-47^, the amine group of 5-HT interacts directly with D399 (Pose_N-D399_) in the lumen-facing open conformation observed in our study. Conversely, in the cytoplasm-facing open conformation in modified VMAT2 with a mutation of Y418S, the amine group of 5-HT points to E312 (Pose_N-E312_)^48^. These observations lead us to speculate on the occurrence of substrate flipping alternation during the substrate transport process. The substrate might rotate in the occluded state during translocation. We extrapolated the potential binding pose of 5-HT in lumen-facing occluded state, using the protein part of TBZ-bound conformation, based on the assumption that the occluded state during transport resembles the occluded conformation triggered by TBZ. This assumption is supported by the FFN206 uptake assay, which showed that R189A and W318A significantly attenuated the transport activity of VMAT2 (Fig. 3F). Ligand-free docking revealed that 5-HT traverses the bipolar cavity and MD simulations showed that 5-HT tends to rotate and point towards D399 in the lumen-facing occluded state (Fig. 3E). Based on the general transport mechanisms of MFS^34,49,50^, we propose a potential substrate flipping mechanism in VMAT2 during transport: in the cytoplasm-facing state during loading, the monoamine substrate associates with E312 through its amine group (Pose_N-E312_); in the lumen-facing occluded state, the substrate might undergo rotation and traverse the bipolar cavity; consequently, the amine group of the substrate points towards D399 (Pose_N-D399_), potentially triggering the conformation transition to lumen-facing state (Fig. 5A).

### MD simulations for protonation and substrate release

As mentioned above, 5-HT stably bound to VMAT2 in the Pose_N-D399_ in three replicas of MD simulations (Fig. 1F). Considering the pH in the presynaptic vesicles is low, this fact prompted us to explore the relevance of the protonation of acidic residues to substrate release. Consequently, we conducted MD simulations to investigate the binding of 5-HT under the protonations of acidic residues. MD simulations indicated that in the lumen-facing configuration, the substrate potentially undergoes a switch from Pose_N-D399_ to Pose_N-E312_ upon protonation of D399, and its release may be initiated when E312 is protonated. The pronation of D426 could potentially trigger substrate release (Fig. 4; Fig. 5A; Table S2). The proton gradient directed toward the cytosol facilitates the transition from the lumen-facing state to the cytoplasm state of VMAT2, suggesting the potential protonations of acidic residues in the translocation pathway, including D33, E312, D399 and D426. However, further studies are needed to fully understand these protonation events on substrate transport of VMAT2.

## Discussion

Our study elucidated the cryo-EM structures of VMAT2 bound to substrates 5-HT/DA, as well as the clinical drugs TBZ/VBZ. Recent investigations have also reported structures of VMAT2 bound to substrate such as 5-HT, or clinical drugs including reserpine/TBZ/ketanserin^45-47,51^. In conjunction with the existing cryo-EM structures of VMAT2-5-HT and our MD simulations studies, we propose a substrate flipping mechanism during transport in VMAT2. This mechanism may involve at least two rounds of substrate flipping: one during substrate loading and translocation accompanied by conformational transitions from a cytoplasm-facing to a lumen-facing state, and the other during substrate release upon protonation of acidic residues (Fig. 5A). It is noteworthy that recent studies have also employed MD simulations on the interplay of protonation and substrate binding of VMAT2, and while their results may vary slightly from ours, likely due to differences in simulation parameters, they consistently highlight the crucial role of D399 in the substrate release process^46,47^.

In addition to the occluded TBZ-bound state and the lumen-facing VBZ-bound state, recent researches have identified an expanded cytoplasm-facing reserpine-bound state and a slightly different lumen-facing ketanserin-bound state^45-47^ (Fig. 5B). These findings suggest a variety of inhibition mechanisms among the drugs targeting VMAT2.

In summary, our study unveils the mechanisms of substrate recognition, translocation, release, and drug inhibition of VMAT2. Our work contributes insights into understanding the mechanisms of monoamine neurotransmitter packing and drug inhibition of VMAT2 and other vesicular monoamine transporters, potentially advancing drug development targeting these transporters for psychiatric disorders.

## Methods

### Expression and purification of BRIL-VMAT2

cDNA of human VMAT2 (UniProt ID: Q05940) gene was synthesized and cloned with N-terminal BRIL sequence into a pCAG vector containing C-terminal tandem His and Flag affinity tag. For the expression of VMAT2 protein, Expi293F cells were used and cultured in SMM 293T-II medium at 37 °C under 5% CO_2_ at 130 rpm in a Multitron-Pro shaker. When the cell density reached approximately 2.0 × 10^6^ cells/mL, Expi293F cells were transiently transfected with the plasmid and polyethylenimines (PEIs) (Polysciences). The total amount of plasmids for transfection of one liter of cell culture is 2 mg and plasmids were pre-mixed with PEIs in 50 mL fresh medium in a 1:2 (w/w) ratio for 20 min before adding the mixture into cell culture.

After transfection for 48 h, cells were harvested by centrifugation at 3000 × rpm and resuspended in the buffer containing 25 mM HEPES pH 7.4, 150 mM NaCl, 1.3 μg/mL aprotinin (Amresco), 0.7 μg/mL pepstatin (Amresco), and 5 μg/mL leupeptin (Amresco). The membrane fraction was extracted with 1% (w/v) n-Dodecyl-β-D-Maltopyranoside (DDM, Anatrace) at 4 °C for 2 h. After extraction, the cell lysate underwent centrifugation at 12,000 × g for 1 h (JA 25.50, Beckman Coulter) and supernatant was loaded to anti-Flag M2 affinity resin (GenScript), washed in buffer containing 25 mM HEPES pH 7.4, 150 mM NaCl, 0.01% LMNG and 0.001% CHS (w/v), and eluted with buffer containing 25 mM HEPES pH 7.4, 150 mM NaCl, 0.01% LMNG and 0.001% CHS (w/v) and 0.4 mg/mL Flag peptide. The elution was then loaded to nickel affinity resin (Ni-NTA, Cytiva), washed with the wash buffer containing 25 mM HEPES pH 7.4, 150 mM NaCl, 0.01% LMNG and 0.001% CHS (w/v), 20 mM imidazole. The protein was eluted with the wash buffer plus 300 mM imidazole. The protein sample after elution was collected and concentrated to 5 mg/mL using a 50 kDa molecular weight cutoff concentrator (Millipore).

### Expression and purification of anti-BRIL Fab

DNA sequences of the heavy and light chains of anti-BRIL Fab were cloned into pCAG expression vectors separately. A His-tag was attached to the C-terminal of the heavy chain. For the expression of Fab, Expi293F cells were cultured and co-transfected by plasmids of heavy chain and light chain under the same conditions as VMAT2. The total amount of heavy chain and light chain plasmids for transfection of one liter of cell culture is 2 mg, 1 mg of each.

Purification of anti-BRIL Fab was conducted as described previously with some modifications^52^. Briefly, after transfection for 48 h, the cell culture media was centrifuged at 3000 × rpm for 10 min, and then the supernatant was loaded to nickel affinity resin, washed with the wash buffer containing 25 mM HEPES pH 7.4, 150 mM NaCl, 50 mM imidazole. The protein was eluted with the wash buffer plus 300 mM imidazole. The protein sample after elution was collected and concentrated to 2 mg/mL using a 50kDa molecular weight cutoff concentrator (Millipore).

### Expression and purification of anti-Fab nanobody

The DNA sequence corresponding to the anti-Fab nanobody (Nb) was inserted into the pET24(+) vector, which features an N-terminal PelB signal sequence^53^ and a C-terminal 6 × His-tag. This construct allowed for the expression and translocation of Nb to the periplasmic space of *Escherichia coli* strain BL21. The bacteria were cultured in LB medium at 37 °C until the optical density at 600 nm (OD_600_) reached approximately 0.8. Subsequently, protein expression was induced by the addition of 1mM isopropyl-β-D-1-thiogalactopyranoside (IPTG) and continued for 16 hours at 16 °C.

For Nb purification, the bacterial cell pellets were lysed by sonication in lysis buffer (25 mM HEPES pH 7.4, 150 mM NaCl, 1 mM PMSF). Cell debris was removed by centrifugation at 16,000 rpm for 1h. Then the supernatant was applied to Ni-NTA, washed with 25 mM HEPES pH 7.4, 150 mM NaCl, 25 mM imidazole and 50 mM imidazole separately. Elution of the protein was performed using an elution buffer containing 25 mM HEPES pH 7.4, 150 mM NaCl and 500 mM imidazole. The protein sample after elution was collected and concentrated to 2 mg/mL using a concentrator with a molecular weight cutoff of 10 kDa (Millipore).

### The BRIL-VMAT2/Fab/Nb complex assembly

For complex assembly, the purified BRIL-VMAT2, anti-BRIL Fab, and anti-Fab Nb were incubated on ice for 1 h at the ratio of 1:1.4:2. Then the samples were applied to a Superose 6 Increase 10/300 GL column (Cytiva) equilibrated with buffer containing 25 mM HEPES pH 7.4, 150 mM NaCl, 0.01% LMNG and 0.001% CHS (w/v) to remove the excess Fab and Nb components. The peak fractions corresponding to the ternary complex were collected and concentrated to 10 mg/mL.

### Cryo-EM sample preparation and data acquisition

The final concentration of protein samples for cryo-EM sample preparation is ∼10 mg/mL. For VMAT2-5-HT/TBZ/VBZ/DA, compounds of serotonin (5-HT, Target-Mol), tetrabenazine (TBZ, Target-Mol), valbenazine (VBZ, MedChemExpress), and dopamine (DA, MedChemExpress) were added to the protein solution at the final concentration of 2.5 mM, 100 μM, 15 μM, and 6.6 mM, individually. 3.5 μL of protein sample with compounds was applied to a glow-discharged Quantifoil R1.2/1.3 400-mesh gold holey carbon grid, blotted for 4.5 s and flash-frozen in liquid ethane cooled by liquid nitrogen with Vitrobot (Mark IV, Thermo Fisher Scientific).

For data acquisition of VMAT2-5-HT/TBZ/VBZ, micrographs were acquired on a Titan Krios microscope (Thermo Fisher Scientific) operated at 300 kV with a Falcon 4i Direct Electron Detector (Thermo Fisher Scientific) and a Selectris X energy filter. EPU was used for automated data collection following standard procedures. A nominal magnification of 130,000 × was used for imaging, yielding a pixel size of 0.932 Å on images. The defocus range was set from −0.9 μm to −1.6 μm. Each micrograph was recorded in a total dose of ∼50 e^–^/Å^2^, with total collection of 13,925, 13,014, and 13,947 movie stacks, respectively, for VMAT2-5-HT/TBZ/VBZ, respectively.

For data acquisition of VMAT2-DA, micrographs were acquired on a Titan Krios microscope (Thermo Fisher Scientific) operated at 300 kV with a K3 Summit direct electron detector (Gatan) and a GIF Quantum energy filter. SerialEM^54^ software was used for automated data collection following standard procedures. A nominal magnification of 105,000 × was used for imaging, yielding a pixel size of 0.832 Å on images. The defocus range was set from −1.3 μm to −1.8 μm. Each micrograph was dose-fractionated to 40 frames in a total dose of ∼50 e^–^/Å^2^, with total collection of 4726 movie stacks for VMAT2-DA.

### Cryo-EM data processing

The data processing pipelines are shown in Figs. S2–S6. Motion correction was performed using Relion v3.1^55^ and CTF estimations were performed using cryoSPARC v3.3.2^56^. For the data processing of VMAT2-5-HT, after several rounds of 2D classification, an initial model with clear TMs was obtained using cryoSPARC. Heterogeneous Refinement, Local refinement and Local CTF refinement yielded a lumen-facing VMAT2-5-HT at the resolution of 2.98 Å. Similarly, we got an occluded VMAT2-TBZ at the resolution of 3.46 Å, a lumen-facing VMAT2-VBZ at the resolution of 3.48 Å, and a lumen-facing VMAT2-DA at the resolution of 3.63 Å.

### Model building

We first built the protein atomic model of VMAT2-5-HT based on the AlphaFold^57,58^ predicted model as a starting template. Using UCSF Chimera^59^ to dock this model into the density maps. Atomic model building based on the density map of VMAT2-5-HT was performed in Coot^60^, and refined in real space using Phenix.real_space_refine.^61^ Validation tools in Phenix^61^ and Molprobity^62^ was used to guide iterative rounds of model adjustment in Coot and refinement in Phenix. For model building of VMAT2-DA/VBZ, atomic model of VMAT2-5-HT was used as initial model, because of the similar conformations. As for VMAT2-TBZ, the AlphaFold predicted model was used as the starting template, and modified subsequently according to the density map.

As regards the modle building of the ligands, we conducted ligand-free docking and calculated the binding free energies through the molecular mechanics-generalized born surface area (MM/GBSA) approach by employing the Schrödinger Maestro program for ligand binding pose determination. Consequently, we constructed the ligand models by referencing to the EM density map and the adjacent environment with Coot. These ligand models underwent further refinement with Phenix.real_space_refine and validated by MD simulations. Visualization of all figures was accomplished using PyMOL (www.pymol.org) or UCSF ChimeraX^63^.

### Transport activity measurements using FFN206

Uptake assay was performed as reported previously^64^, with some modifications. Briefly, HEK293T cells were cultured in 6-well plates until reaching approximately 50-60% confluence, and expression plasmid pCAG-WT or pCAG-mutants were transfected with PEIs following the manufacturer’s instructions. After transfection for 48 h, the medium was discarded, and cells were incubated with 2 mL of assay buffer (125 mM sodium gluconate, 4.8 mM potassium gluconate, 1.2 mM NaH_2_PO_4_, 1.2 mM MgSO_4_, 1.3 mM CaCl_2_, 5.6 mM glucose and 25 mM HEPES pH 7.4) at 37 °C for 15 min. Subsequently, the fluorescent indicator substrate FFN206 (Target-Mol) was added to the cells at a final concentration of 5 μM in assay buffer, and the cells were incubated at 37 °C for 1 h. Uptake was terminated by washing cells three times with 2 mL ice-cold phosphate-buffered saline (PBS). Cells were lysed with PBS containing 1% TritonX-100, and the fluorescence intensity of FFN206 was measured with excitation/emission wavelength at 360 nm/460 nm using a microplate reader (BioTek Synergy2 Microplate Reader). All statistical analyses were performed with one-way ANOVA followed by Dunnett’s multiple comparisons test by comparing WT with mutants using GraphPad Prism v9.3.1. Transport assays were biologically repeated at least three times. The average fluorescent signals of untransfected cells were treated as background correction and the average fluorescent signals of VMAT2_WT_ were defined as 100% uptake.

For the transport inhibiton assay, HEK293T cells were cultured in 6-well plates, until reaching approximately 50-60% confluence, expression plasmid pCAG-VMAT2_WT_ or pCAG-VMAT2_EM_ were transfected with Lipofectamine^TM^ 2000 according to the manufacturer’s instructions. After transfection for 24 h, the medium was discarded and cells were divided into 24-well plates. 10 mM sodium butyrate was added, and cells were cultivated at 30 °C for 24 h. Afterward, cells were treated with assay buffer containing 10 mM Glyco-diosgenin (GDN) with inhibitors (TBZ or VBZ) or DMSO (as control), and incubated at 37 °C for 15 min. Next, cells were incubated for another 1 h in uptake buffer (assay buffer containing FFN206, final concentration 5μM) at 37 °C and washed three times with 2 mL ice-cold PBS. Cells were lysed using PBS containing 1% TritonX-100 and transformed to a 96-well plate, the fluorescence intensity of FFN206 was measured with excitation/emission wavelength at 360 nm/460 nm using a microplate reader (BioTek Synergy2 Microplate Reader). In this assay, 5 μM TBZ and 5 μM VBZ were applied as inhibitors. The average fluorescent signals of untransfected cells were treated as background correction and the average fluorescent signals of VMAT2_WT_ were defined as 100% uptake.

### Cell surface biotinylation and immunoblot analysis

Cell surface biotinylation and immunoblot analysis were performed as reported previously with some modifications^65^. Briefly, HEK293T cells were cultured in 6-well plates until reaching approximately 70% confluence, and expression plasmid pCAG-WT or pCAG-mutants were transfected with PEIs following the manufacturer’s instructions. After 48 h, The cell culture medium was removed and cells were suspended with 1 mL ice-cold PBS and then treated with 1 mL sulfo-N-hydroxysuccinimide-SS-biotin (Thermo Scientific) (1 mg/mL in PBS) at room temperature for 1 h. Afterward, cells were washed three times with 1 mL ice-cold PBS containing 100 mM glycine and incubated at 4°C for 10 min in the same buffer. Then, cells were lysed with 500 μL lysis buffer (25 mM Tris pH 7.4, 150 mM NaCl, 1 mM EDTA, 0.1% SDS, and 1% Triton X-100, containing protease inhibitors) at 4°C for 2 h with shaking. After centrifugation at 13,000 g for 10 min, the supernatants were incubated with 50 μL of streptavidin-agarose beads (Thermo Scientific) at room temperature for 1 h. The beads were then centrifuged at 500 g for 1 min, washed three times with ice-cold lysis buffer, and incubated with 100 μL of 2 × Laemmli buffer containing 100 mM dithiothreitol at room temperature for 1 h to recover the cell surface proteins. Samples were separated using SDS-polyacrylamide gel electrophoresis followed by western-blot analysis. All proteins were detected using anti-Flag-HRP antibody (GNI) (1: 5000 dilution). Then the same membrane was stripped at room temperature for 1 h. Na^+^/K^+^-ATPase was used as loading control for normalization and was detected with rabbit anti-Na^+^/K^+^-ATPase monoclonal antibody (1:10,000 dilution) (ABclonal) combined with secondary HRP goat anti-rabbit monoclonal antibody (1:10,000 dilution) (ABclonal).

### Molecular dynamics simulations

Molecular dynamics (MD) simulations were conducted for VMAT2-5-HT/DA/TBZ/VBZ. The initial proteins include residues 18 to 46 and 117 to 475 for VMAT2-5-HT/VBZ, residues 18 to 46 and 132 to 475 for VMAT2-DA and residues 16 to 50, 117 to 318 and 324 to 476 for VMAT2-TBZ. Charged residues were protonated assuming pH 7.4 unless otherwise specified, and all the N- and C-termini of proteins were capped with acetyl and N-methyl groups, respectively. To build the system, protein-ligand complex was firstly embedded in a POPC bilayer (∼302 molecules) and then solvated in a water box (117 × 117 × 90 Å^3^). Finally, 150 mM NaCl was added to neutralize the system. Details of system composition are listed in Table S2.

MD simulations were performed with GROMACS 2021.4^66^. CHARMM36 force field was used for protein, POPC lipid and ions^67,68^, and TIP3P model^69^ was used for water. The force fields of four ligands (5-HT/DA/TBZ/VBZ) were generated by CgenFF program^70,71^. After energy minimization by steepest descent algorithm, the systems were simulated in NVT ensemble to relax lipid tail, lipid head and solvent (water and ions) in steps for 1.5 ns. Then, the systems were further relaxed in NPT ensemble. Constraints on 1) lipids and solvent, 2) protein sidechain, 3) ligand and 4) protein Cα atoms were gradually released each for 2 ns. After pre-equilibrations, 500 or 600 ns production simulations were run without any constraint. Periodic boundary condition was applied, and the temperature and pressure were kept at 310 K and 1 atm by Nose-Hoover thermostat^72,73^ and Parrinello-Rahman barostat^74^, respectively. Long range electrostatic interactions were described using the particle mesh Ewald (PME) method^75^ with a cutoff of 12 Å. Van der Waals interactions were smoothly switched to zero from 10 to 12 Å. All bonds involving hydrogen atoms were constrained by the LINCS algorithm^76^ to allow a time step of 2 fs. Each system was run for three replicas with different initial velocities and the total simulation time was 18.3 µs.

## Data availability

Atomic coordinates of VMAT2-5-HT/DA/TBZ/VBZ have been deposited in the Protein Data Bank (http://www.rcsb.org) under the accession codes 8X3K, 8X4V, 8X4U, and 8XT6, respectively. The corresponding electron microscopy maps have been deposited in the Electron Microscopy Data Bank (https://www.ebi.ac.uk/pdbe/emdb/), under the accession codes EMD-38038, EMD-38052, EMD-38051, and EMD-38640, respectively.

## Acknowledgments

We thank the Cryo-EM Center of Institutes of Biomedical Sciences of Fudan University for providing the cryo-EM facility support and the computational facility support. We thank Prof. Yanhui Xu, Prof. Zhenguo Chen, Hui Zhao, Yulei Ren and Anqi Dong for technical support in the Cryo-EM Center. We thank Prof. Zhe Zhang, from Peking University, for kindly providing the plasmids of anti-BRIL Fab and nanobody. This work was funded by Shanghai Rising-Star Program (21QA1401200 to Y.Z.), the National Natural Science Foundation of China (32171196 to Y.Z.), National Key R&D Program of China (2021YFA1300101 to Y.Z.), Start-up funds from Fudan University and Shanghai Fifth People’s Hospital, and Shenzhen Natural Science Foundation Project (GXWD20201231105722002-20200831175432002 to H.L.).

## Author contributions

Y.Z. conceived and designed the project. F.W.and J.W. prepared protein purification. W.Z. performed functional transport assay. F.W. performed cryo-EM data acquisition. Y.Z. and F.W. contributed to the cryo-EM data processing and model building. H.L. performed MD simulations. Y.Z. wrote the manuscript. All authors analyzed the data and contributed to manuscript preparation.

## References

1. Eiden, L.E., Schäfer, M.K.H., Weihe, E. & Schütz, B. The vesicular amine transporter family (SLC18): amine/proton antiporters required for vesicular accumulation and regulated exocytotic secretion of monoamines and acetylcholine. Pflugers Archiv : European Journal of Physiology 447, 636–640 (2004).

2. Nirenberg, M.J., Liu, Y., Peter, D., Edwards, R.H. & Pickel, V.M. The vesicular monoamine transporter 2 is present in small synaptic vesicles and preferentially localizes to large dense core vesicles in rat solitary tract nuclei. Proc Natl Acad Sci U S A 92, 8773–7 (1995).

3. Alwindi, M. & Bizanti, A. Vesicular monoamine transporter (VMAT) regional expression and roles in pathological conditions. Heliyon 9, e22413 (2023).

4. Anne, C. & Gasnier, B. Vesicular neurotransmitter transporters: mechanistic aspects. Curr Top Membr 73, 149–74 (2014).

5. Arvidsson, U., Riedl, M., Elde, R. & Meister, B. Vesicular acetylcholine transporter (VAChT) protein: a novel and unique marker for cholinergic neurons in the central and peripheral nervous systems. J Comp Neurol 378, 454–67 (1997).

6. Hiasa, M. et al. Identification of a mammalian vesicular polyamine transporter. Sci Rep 4, 6836 (2014).

7. Moriyama, Y., Hatano, R., Moriyama, S. & Uehara, S. Vesicular polyamine transporter as a novel player in amine-mediated chemical transmission. Biochim Biophys Acta Biomembr 1862, 183208 (2020).

8. Lawal, H.O. & Krantz, D.E. SLC18: Vesicular neurotransmitter transporters for monoamines and acetylcholine. Molecular Aspects of Medicine 34, 360–372 (2013).

9. Schuldiner, S., Shirvan, A. & Linial, M. Vesicular neurotransmitter transporters: from bacteria to humans. Physiological Reviews 75, 369–392 (1995).

10. Bernstein, A.I., Stout, K.A. & Miller, G.W. The vesicular monoamine transporter 2: an underexplored pharmacological target. Neurochemistry International 73, 89–97 (2014).

11. Eiden, L.E. & Weihe, E. VMAT2: a dynamic regulator of brain monoaminergic neuronal function interacting with drugs of abuse. Annals of the New York Academy of Sciences 1216, 86–98 (2011).

12. Erickson, J.D., Schafer, M.K., Bonner, T.I., Eiden, L.E. & Weihe, E. Distinct pharmacological properties and distribution in neurons and endocrine cells of two isoforms of the human vesicular monoamine transporter. Proceedings of the National Academy of Sciences of the United States of America 93, 5166–5171 (1996).

13. Weihe, E., Schäfer, M.K., Erickson, J.D. & Eiden, L.E. Localization of vesicular monoamine transporter isoforms (VMAT1 and VMAT2) to endocrine cells and neurons in rat. Journal of Molecular Neuroscience : MN 5, 149–164 (1994).

14. Teleanu, R.I. et al. Neurotransmitters-Key Factors in Neurological and Neurodegenerative Disorders of the Central Nervous System. International Journal of Molecular Sciences 23(2022).

15. Isingrini, E. et al. Selective genetic disruption of dopaminergic, serotonergic and noradrenergic neurotransmission: insights into motor, emotional and addictive behaviour. Journal of Psychiatry & Neuroscience : JPN 41, 169–181 (2016).

16. Mulvihill, K.G. Presynaptic regulation of dopamine release: Role of the DAT and VMAT2 transporters. Neurochemistry International 122(2019).

17. Erickson, J.D. & Eiden, L.E. Functional identification and molecular cloning of a human brain vesicle monoamine transporter. J Neurochem 61, 2314–7 (1993).

18. Coppen, A. The biochemistry of affective disorders. The British Journal of Psychiatry : the Journal of Mental Science 113, 1237–1264 (1967).

19. Edwards, R.H. The neurotransmitter cycle and quantal size. Neuron 55, 835–858 (2007).

20. Guillot, T.S. & Miller, G.W. Protective actions of the vesicular monoamine transporter 2 (VMAT2) in monoaminergic neurons. Molecular Neurobiology 39, 149–170 (2009).

21. Lohr, K.M. et al. Increased vesicular monoamine transporter enhances dopamine release and opposes Parkinson disease-related neurodegeneration in vivo. Proceedings of the National Academy of Sciences of the United States of America 111, 9977–9982 (2014).

22. Chen, H.-H. et al. Therapeutic effects of honokiol on motor impairment in hemiparkinsonian mice are associated with reversing neurodegeneration and targeting PPARγ regulation. Biomedicine & Pharmacotherapy = Biomedecine & Pharmacotherapie 108, 254–262 (2018).

23. Taylor, S.F., Koeppe, R.A., Tandon, R., Zubieta, J.K. & Frey, K.A. In vivo measurement of the vesicular monoamine transporter in schizophrenia. Neuropsychopharmacology : Official Publication of the American College of Neuropsychopharmacology 23, 667–675 (2000).

24. Chaudhry, F.A., Boulland, J.-L., Jenstad, M., Bredahl, M.K.L. & Edwards, R.H. Pharmacology of neurotransmitter transport into secretory vesicles. Handbook of Experimental Pharmacology (2008).

25. Wimalasena, K. Vesicular monoamine transporters: structure-function, pharmacology, and medicinal chemistry. Medicinal Research Reviews 31, 483–519 (2011).

26. Tarakad, A. & Jimenez-Shahed, J. VMAT2 Inhibitors in Neuropsychiatric Disorders. CNS Drugs 32, 1131–1144 (2018).

27. Kaur, N., Kumar, P., Jamwal, S., Deshmukh, R. & Gauttam, V. Tetrabenazine: Spotlight on Drug Review. Annals of Neurosciences 23, 176–185 (2016).

28. Canney, D.J., Kung, M.P. & Kung, H.F. Amino- and amido-tetrabenazine derivatives: synthesis and evaluation as potential ligands for the vesicular monoamine transporter. Nuclear Medicine and Biology 22, 527–535 (1995).

29. Kim, E.S. Valbenazine: First Global Approval. Drugs 77, 1123–1129 (2017).

30. Davis, M.C., Miller, B.J., Kalsi, J.K., Birkner, T. & Mathis, M.V. Efficient Trial Design - FDA Approval of Valbenazine for Tardive Dyskinesia. The New England Journal of Medicine 376, 2503–2506 (2017).

31. Njus, D., Kelley, P.M. & Harnadek, G.J. Bioenergetics of secretory vesicles. Biochimica Et Biophysica Acta 853, 237–265 (1986).

32. Omote, H., Miyaji, T., Juge, N. & Moriyama, Y. Vesicular neurotransmitter transporter: bioenergetics and regulation of glutamate transport. Biochemistry 50, 5558–5565 (2011).

33. Mukherjee, S. et al. Synthetic antibodies against BRIL as universal fiducial marks for single-particle cryoEM structure determination of membrane proteins. Nature Communications 11, 1598 (2020).

34. Drew, D., North, R.A., Nagarathinam, K. & Tanabe, M. Structures and General Transport Mechanisms by the Major Facilitator Superfamily (MFS). Chemical Reviews 121, 5289–5335 (2021).

35. Drew, D. & Boudker, O. Shared Molecular Mechanisms of Membrane Transporters. Annual Review of Biochemistry 85, 543–572 (2016).

36. Frank, S. et al. Effect of Deutetrabenazine on Chorea Among Patients With Huntington Disease: A Randomized Clinical Trial. JAMA 316, 40–50 (2016).

37. Wang, H., Chen, X., Li, Y., Tang, T.-S. & Bezprozvanny, I. Tetrabenazine is neuroprotective in Huntington’s disease mice. Molecular Neurodegeneration 5, 18 (2010).

38. Yohn, S.E. et al. Not All Antidepressants Are Created Equal: Differential Effects of Monoamine Uptake Inhibitors on Effort-Related Choice Behavior. Neuropsychopharmacology : Official Publication of the American College of Neuropsychopharmacology 41, 686–694 (2016).

39. Yaffe, D., Radestock, S., Shuster, Y., Forrest, L.R. & Schuldiner, S. Identification of molecular hinge points mediating alternating access in the vesicular monoamine transporter VMAT2. Proceedings of the National Academy of Sciences of the United States of America 110, E1332–E1341 (2013).

40. Peter, D., Jimenez, J., Liu, Y., Kim, J. & Edwards, R.H. The chromaffin granule and synaptic vesicle amine transporters differ in substrate recognition and sensitivity to inhibitors. The Journal of Biological Chemistry 269, 7231–7237 (1994).

41. Grigoriadis, D.E., Smith, E., Hoare, S.R.J., Madan, A. & Bozigian, H. Pharmacologic Characterization of Valbenazine (NBI-98854) and Its Metabolites. The Journal of Pharmacology and Experimental Therapeutics 361, 454–461 (2017).

42. Kiyoi, T. et al. High-throughput screening system for dynamic monitoring of exocytotic vesicle trafficking in mast cells. PloS One 13, e0198785 (2018).

43. Ugolev, Y., Segal, T., Yaffe, D., Gros, Y. & Schuldiner, S. Identification of conformationally sensitive residues essential for inhibition of vesicular monoamine transport by the noncompetitive inhibitor tetrabenazine. The Journal of Biological Chemistry 288, 32160–32171 (2013).

44. Yaffe, D., Vergara-Jaque, A., Forrest, L.R. & Schuldiner, S. Emulating proton-induced conformational changes in the vesicular monoamine transporter VMAT2 by mutagenesis. Proceedings of the National Academy of Sciences of the United States of America 113, E7390–E7398 (2016).

45. Pidathala, S. et al. Mechanisms of neurotransmitter transport and drug inhibition in human

46. Wang, Y. et al. Transport and inhibition mechanism for VMAT2-mediated synaptic vesicle loading of monoamines. Cell Research 34, 47–57 (2024).

47. Wu, D. et al. Transport and inhibition mechanisms of human VMAT2. Nature 626, 427–434 (2024).

48. Pidathala, S. et al. Mechanisms of neurotransmitter transport and drug inhibition in human VMAT2. Nature 623, 1086–1092 (2023).

49. Quistgaard, E.M., Löw, C., Guettou, F. & Nordlund, P. Understanding transport by the major facilitator superfamily (MFS): structures pave the way. Nature Reviews. Molecular Cell Biology 17, 123–132 (2016).

50. Yan, N. Structural Biology of the Major Facilitator Superfamily Transporters. Annual Review of Biophysics 44, 257–283 (2015).

51. Dalton, M.P., Cheng, M.H., Bahar, I. & Coleman, J.A. Structural mechanisms for VMAT2 inhibition by tetrabenazine. eLife (2023).

52. Dang, Y. et al. Molecular mechanism of substrate recognition by folate transporter SLC19A1. Cell Discovery 8, 141 (2022).

53. Steiner, D., Forrer, P., Stumpp, M.T. & Plückthun, A. Signal sequences directing cotranslational translocation expand the range of proteins amenable to phage display. Nature Biotechnology 24, 823–831 (2006).

54. Mastronarde, D.N. Automated electron microscope tomography using robust prediction of specimen movements. Journal of structural biology 152, 36–51 (2005).

55. He, S. & Scheres, S.H. Helical reconstruction in RELION. Journal of structural biology 198, 163–176 (2017).

56. Punjani, A., Rubinstein, J.L., Fleet, D.J. & Brubaker, M.A. cryoSPARC: algorithms for rapid unsupervised cryo-EM structure determination. Nature methods 14, 290–296 (2017).

57. Jumper, J. et al. Highly accurate protein structure prediction with AlphaFold. Nature 596, 583–589 (2021).

58. Varadi, M. et al. AlphaFold Protein Structure Database: massively expanding the structural coverage of protein-sequence space with high-accuracy models. Nucleic acids research 50, D439–D444 (2022).

59. Pettersen, E.F. et al. UCSF Chimera—a visualization system for exploratory research and analysis. Journal of computational chemistry 25, 1605–1612 (2004).

60. Emsley, P., Lohkamp, B., Scott, W.G. & Cowtan, K. Features and development of Coot. Acta Crystallographica Section D: Biological Crystallography 66, 486–501 (2010).

61. Adams, P.D. et al. PHENIX: a comprehensive Python-based system for macromolecular structure solution. Acta Crystallographica Section D: Biological Crystallography 66, 213–221 (2010).

62. Chen, V.B. et al. MolProbity: all-atom structure validation for macromolecular crystallography. Acta Crystallographica Section D: Biological Crystallography 66, 12–21 (2010).

63. Pettersen, E.F. et al. UCSF ChimeraX: Structure visualization for researchers, educators, and developers. Protein Science 30, 70–82 (2021).

64. Pidathala, S. et al. Mechanisms of neurotransmitter transport and drug inhibition in human

65. Shan, Z. et al. Cryo-EM structures of human organic anion transporting polypeptide OATP1B1. Cell Res 33, 940–951 (2023).

66. Abraham, M.J. et al. GROMACS: High performance molecular simulations through multi-level parallelism from laptops to supercomputers. SoftwareX 1-2, 19-25 (2015).

67. Huang, J. et al. CHARMM36m: an improved force field for folded and intrinsically disordered proteins. Nature Methods 14, 71–73 (2017).

68. Klauda, J.B. et al. Update of the CHARMM all-atom additive force field for lipids: validation on six lipid types. The Journal of Physical Chemistry. B 114, 7830–7843 (2010).

69. Mark, P. & Nilsson, L. Structure and Dynamics of the TIP3P, SPC, and SPC/E Water Models at 298 K. The Journal of Physical Chemistry A 105, 9954-9960 (2001).

70. Vanommeslaeghe, K. & MacKerell, A.D., Jr. Automation of the CHARMM General Force Field (CGenFF) I: Bond Perception and Atom Typing. Journal of Chemical Information and Modeling 52, 3144–3154 (2012).

71. Vanommeslaeghe, K., Raman, E.P. & MacKerell, A.D., Jr. Automation of the CHARMM General Force Field (CGenFF) II: Assignment of Bonded Parameters and Partial Atomic Charges. Journal of Chemical Information and Modeling 52, 3155-3168 (2012).

72. Hoover, W.G. Canonical dynamics: Equilibrium phase-space distributions. Physical Review A 31, 1695–1697 (1985).

73. Nosé, S. A unified formulation of the constant temperature molecular dynamicss methods. The Journal of Chemical Physics 81, 511–519 (1984).

74. Parrinello, M. & Rahman, A. Polymorphic transitions in single crystals: A new molecular dynamicss method. Journal of Applied Physics 52, 7182–7190 (1981).

75. Essmann, U. et al. A smooth particle mesh Ewald method. The Journal of Chemical Physics 103, 8577–8593 (1995).

76. Hess, B. P-LINCS: A Parallel Linear Constraint Solver for Molecular Simulation. Journal of Chemical Theory and Computation 4, 116–122 (2008).

