## Supplementary Material for "Drug inhibition and substrate alternating flipping mechanisms of human VMAT2"

**This PDF file includes:**

Figures S1 to S8

Tables S1 to S2

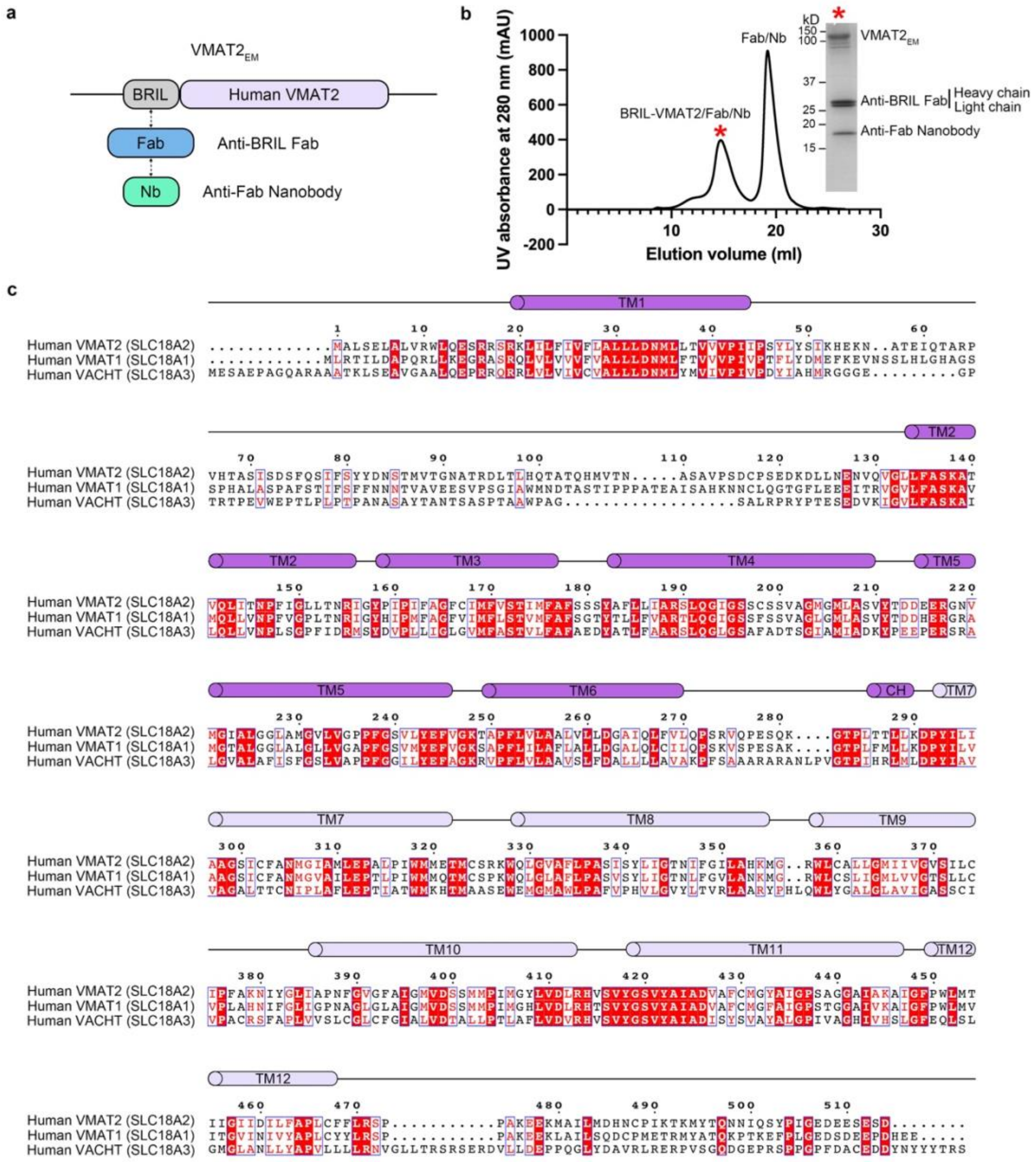

**Fig. S1 | Cryo-EM sample design and sequence alignment of VMAT2.**

**A** Schematic diagram of the $VMAT2_{EM}$ construct. **B** Size exclusion chromatography (SEC) of the VMAT2 by Superose 6 and the protein peak detected by SDS-PAGE gel. **C** Sequence alignment among human SLC18 family transporters.

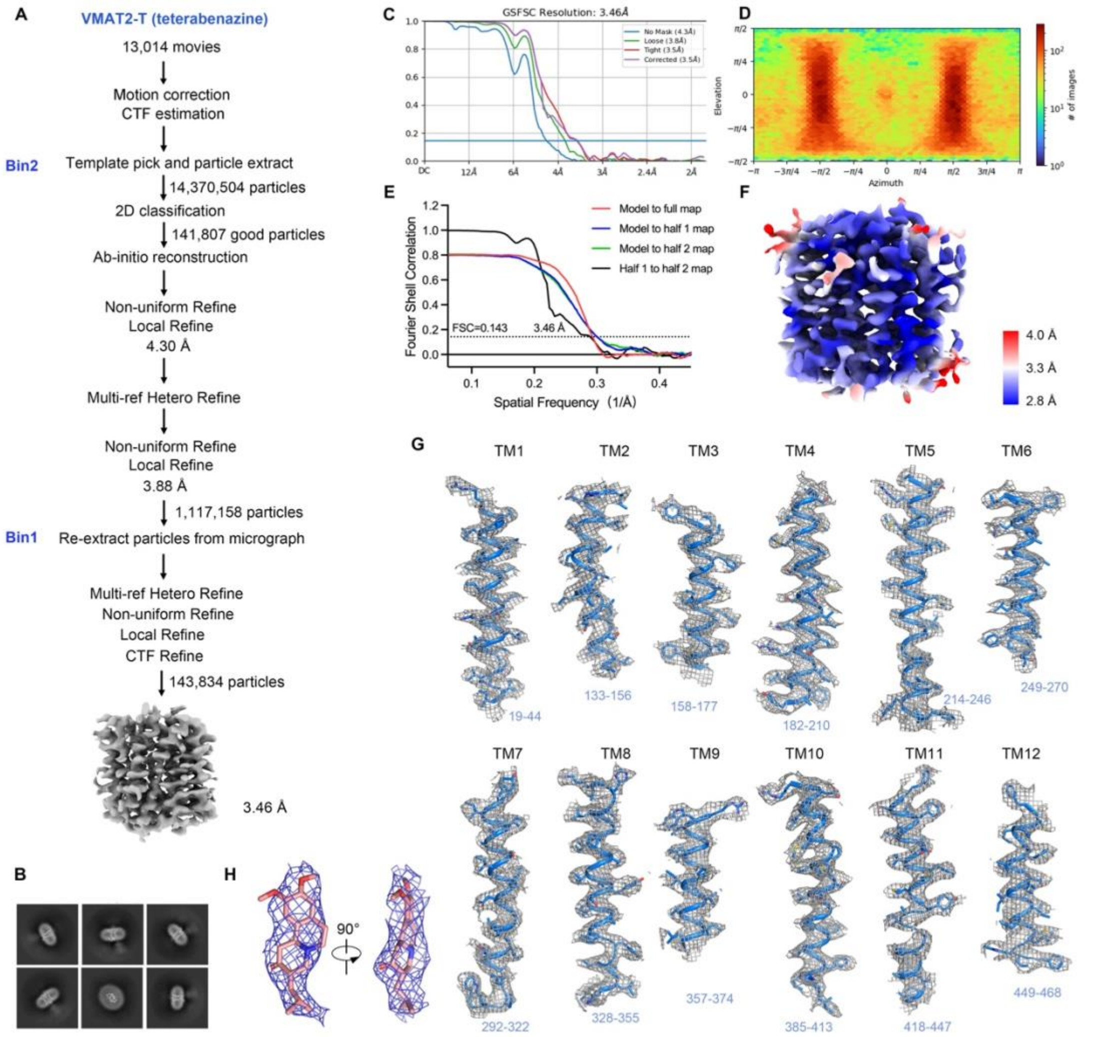

**Fig. S2 | Cryo-EM data processing of VMAT2 bound to tetrabenzazine.**

**A** Flowchart of cryo-EM data processing of VMAT2-TBZ. TBZ, tetrabenazine. **B** Representative 2D class averages. **C** Fourier shell correlation (FSC) curves between two half maps generated by cryoSPARC. **D** Angular distribution of particles contributing to the final cryo-EM map of VMAT2-TBZ. **E** Gold-standard FSC curve between two half maps (black) with indicated resolution at 0.143, and FSC curves between the atomic model refined against full map (red) and half 1 map (blue) or half 2 map (green). **F** The local resolution map of VMAT2-TBZ. **G** The cryo-EM density maps of all transmembrane helices are shown as mesh (5σ) colored in gray, with atomic models shown as cartoon and sidechains shown as sticks, and colored in marine. **H** The cryo-EM density map of TBZ is shown as mesh (5σ), and colored in salmon.

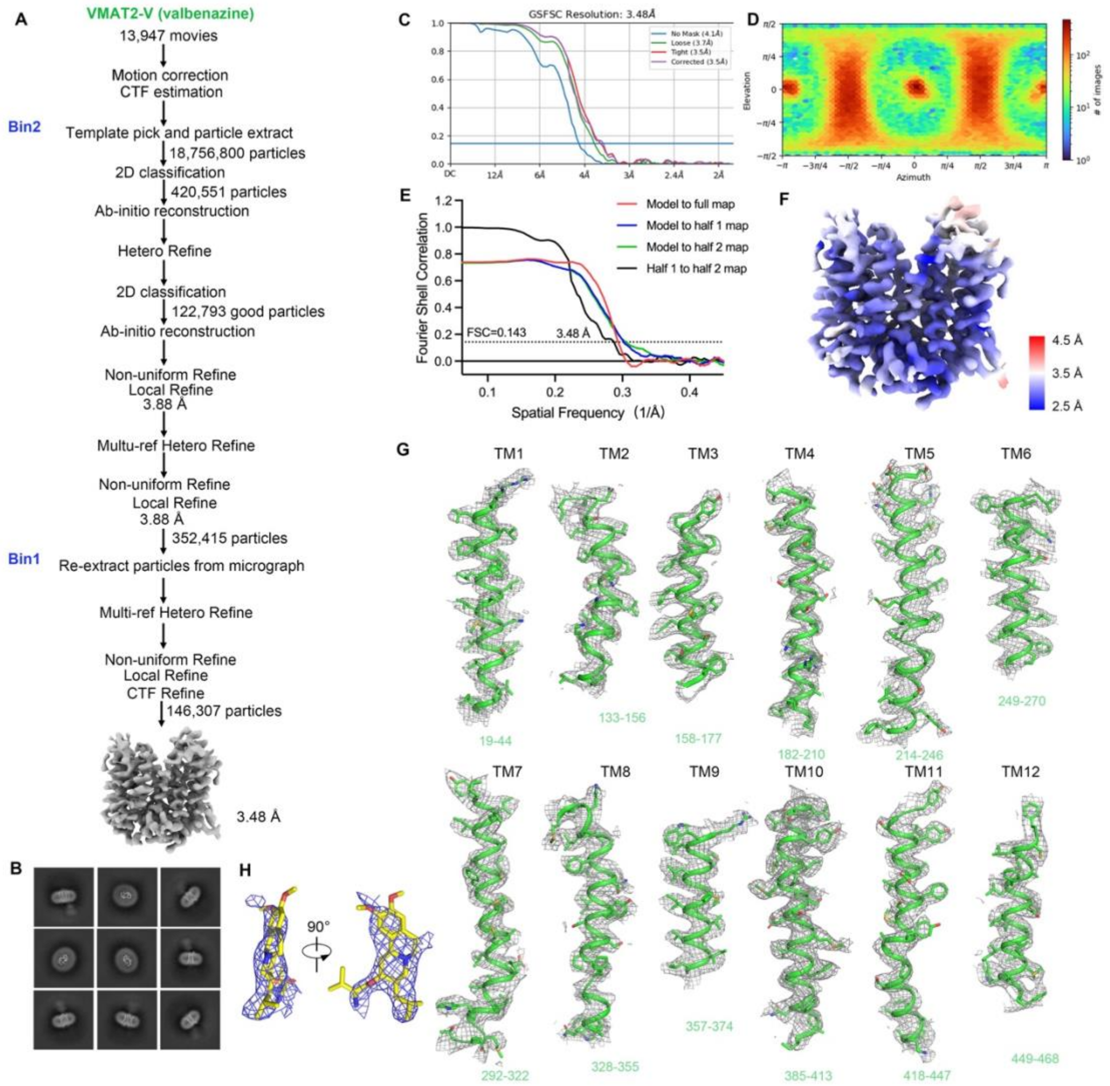

**Fig. S3 | Cryo-EM data processing of VMAT2 bound to valbenazine.**
**A** Flowchart of cryo-EM data processing of VMAT2-VBZ. VBZ, valbenazine. **B** Representative 2D class averages. **C** Fourier shell correlation (FSC) curves between two half maps generated by cryoSPARC. **D** Angular distribution of particles contributing to the final cryo-EM map of VMAT2-VBZ. **E** Gold-standard FSC curve between two half maps (black) with indicated resolution at 0.143, and FSC curves between the atomic model refined against full map (red) and half 1 map (blue) or half 2 map (green). **F** The local resolution map of VMAT2-VBZ. **G** The cryo-EM density maps of all transmembrane helices are shown as mesh (5σ) colored in gray, with atomic models shown as cartoon and sidechains shown as sticks, and colored in green. **H** The cryo-EM density map of VBZ is shown as mesh (5σ), and colored in yellow.

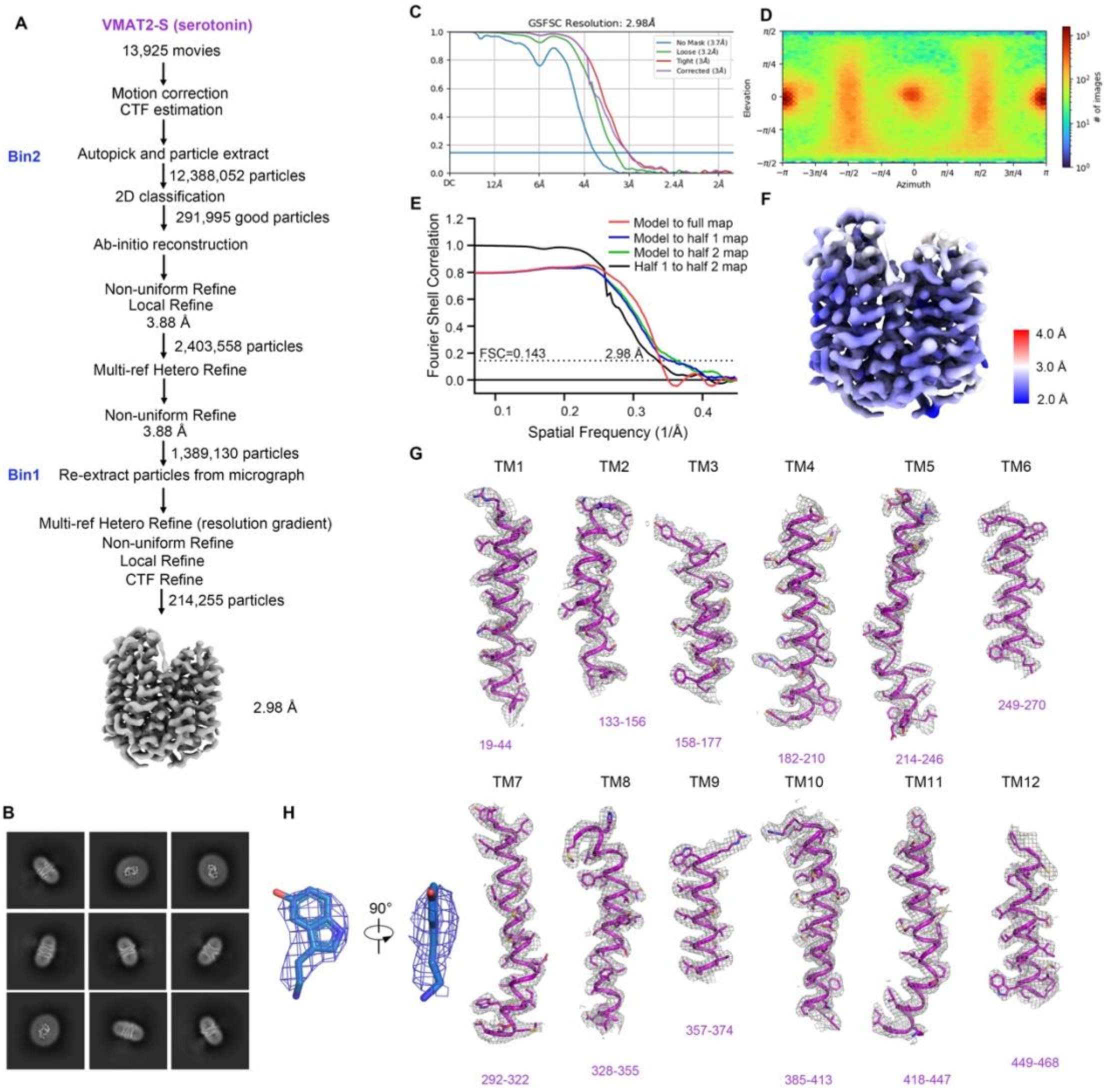

**Fig. S4 | Cryo-EM data processing of VMAT2 bound to serotonin.**

**A** Flowchart of cryo-EM data processing of VMAT2-5-HT. 5-HT, serotonin. **B** Representative 2D class averages. **C** Fourier shell correlation (FSC) curves between two half maps generated by cryoSPARC. **D** Angular distribution of particles contributing to the final cryo-EM map of VMAT2-5-HT. **E** Gold-standard FSC curve between two half maps (black) with indicated resolution at 0.143, and FSC curves between the atomic model refined against full map (red) and half 1 map (blue) or half 2 map (green). **F** The local resolution map of VMAT2-5-HT. **G** The cryo-EM density maps of all transmembrane helices are shown as mesh (5σ) colored in gray, with atomic models shown as cartoon and sidechains shown as sticks, and colored in purple. **H** The cryo-EM density map of 5-HT is shown as mesh (5σ), and colored in marine.

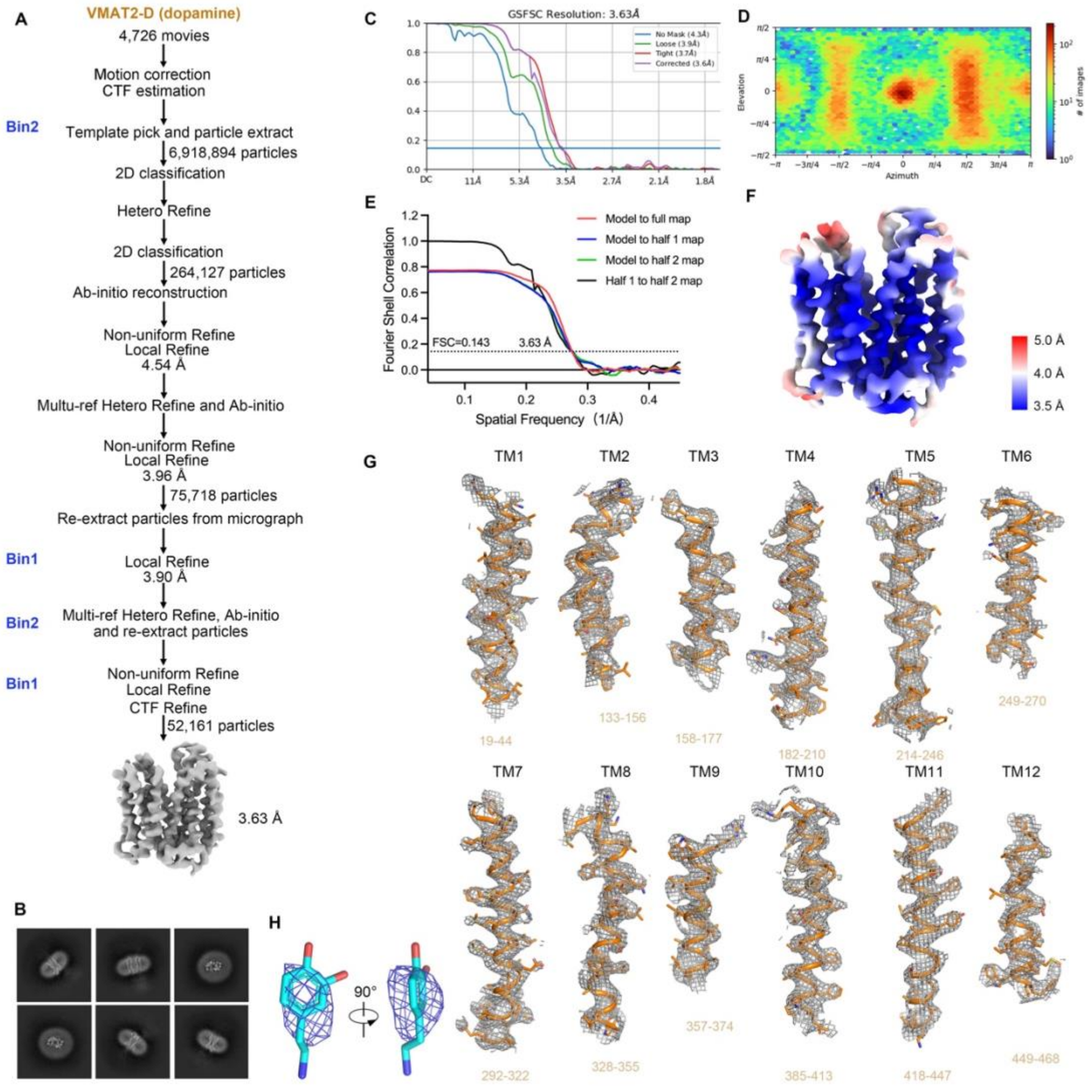

**Fig. S5 | Cryo-EM data processing of VMAT2 bound to dopamine.**

**A** Flowchart of cryo-EM data processing of VMAT2-DA. DA, dopamine. **B** Representative 2D class averages. **C** Fourier shell correlation (FSC) curves between two half maps generated by cryoSPARC. **D** Angular distribution of particles contributing to the final cryo-EM map of VMAT2-DA. **E** Gold-standard FSC curve between two half maps (black) with indicated resolution at 0.143, and FSC curves between the atomic model refined against full map (red) and half 1 map (blue) or half 2 map (green). **F** The local resolution map of VMAT2-DA. **G** The cryo-EM density maps of all transmembrane helices are shown as mesh (5σ) colored in gray, with atomic models shown as cartoon and sidechains shown as sticks, and colored in orange. **H** The cryo-EM density map of DA is shown as mesh (5σ), and colored in cyan.

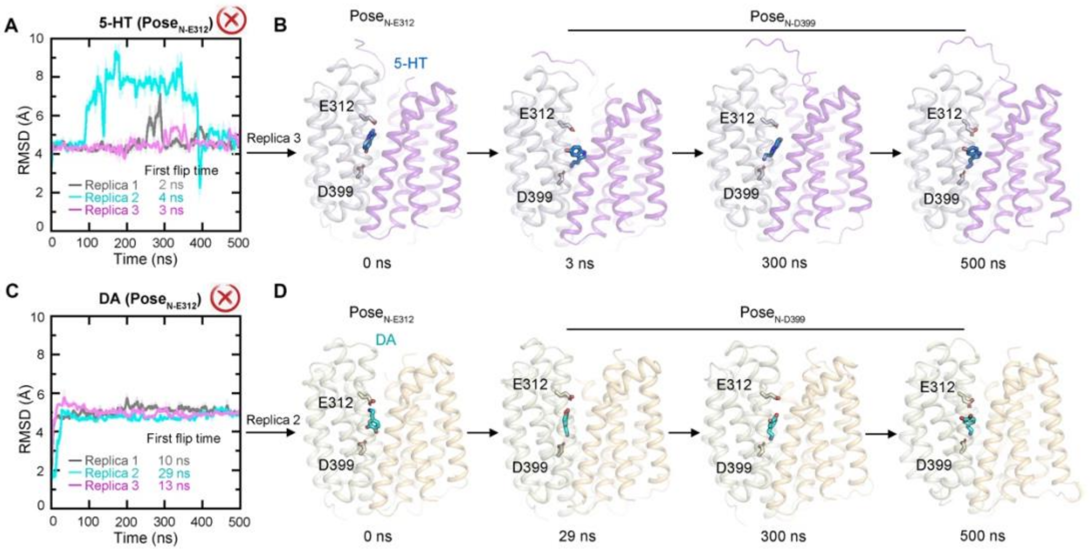

**Fig. S6 | Molecular dynamics (MD) simulations of VMAT2 structures.**
**A** Root mean square deviation (RMSD) plots of MD simulations for 5-HT (Pose$_{\text{N-E312}}$). **B** MD snapshots of the 5-HT (Pose$_{\text{N-E312}}$) bound lumen-facing VMAT2 structure. Frames are selected as shown. 5-HT would flip from Pose$_{\text{N-E312}}$ to Pose$_{\text{N_D399}}$. **C** RMSD plots of MD simulations for DA (Pose$_{\text{N-E312}}$). **D** MD snapshots of the DA (Pose$_{\text{N-E312}}$) bound lumen-facing VMAT2 structure. Frames are selected as shown. DA would flip from Pose$_{\text{N-E312}}$ to Pose$_{\text{N_D399}}$. 5-HT, serotonin; DA, dopamine.

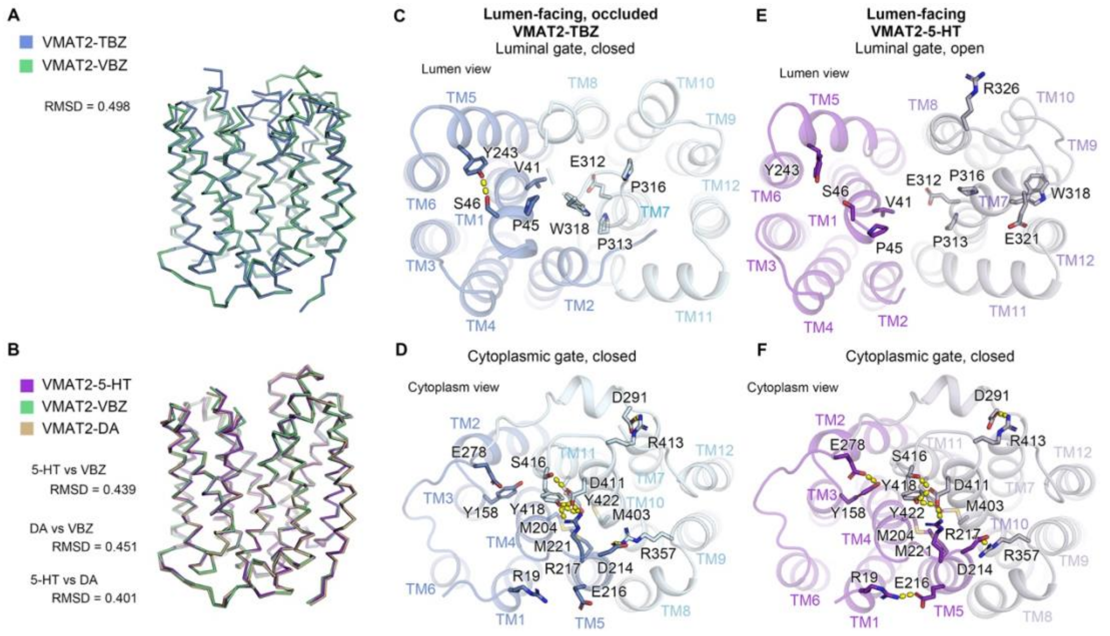

**Fig. S7 | The luminal and cytoplasmic gates in different conformations of VMAT2**

**A** Superposition of VMAT2-TBZ (lumen-facing occluded, blue) and VMAT2-VBZ (lumen-facing open, green) shown in ribbon. **B** Superposition of VMAT2-5-HT, VMAT2-VBZ and VMAT2-DA, all three are lumen-facing open conformations. RMSDs show only minor differences between them. **C** The luminal gate in the VMAT2-TBZ from the lumen view. **D** The cytoplasmic gate in the VMAT2-TBZ from the cytoplasm view. **E** The luminal gate in the VMAT2-5-HT from the lumen view. **F** The cytoplasmic gate in the VMAT2-5-HT from the cytoplasm view. TBZ, tetrabenazine; VBZ, valbenazine; 5-HT, serotonin; DA, dopamine.

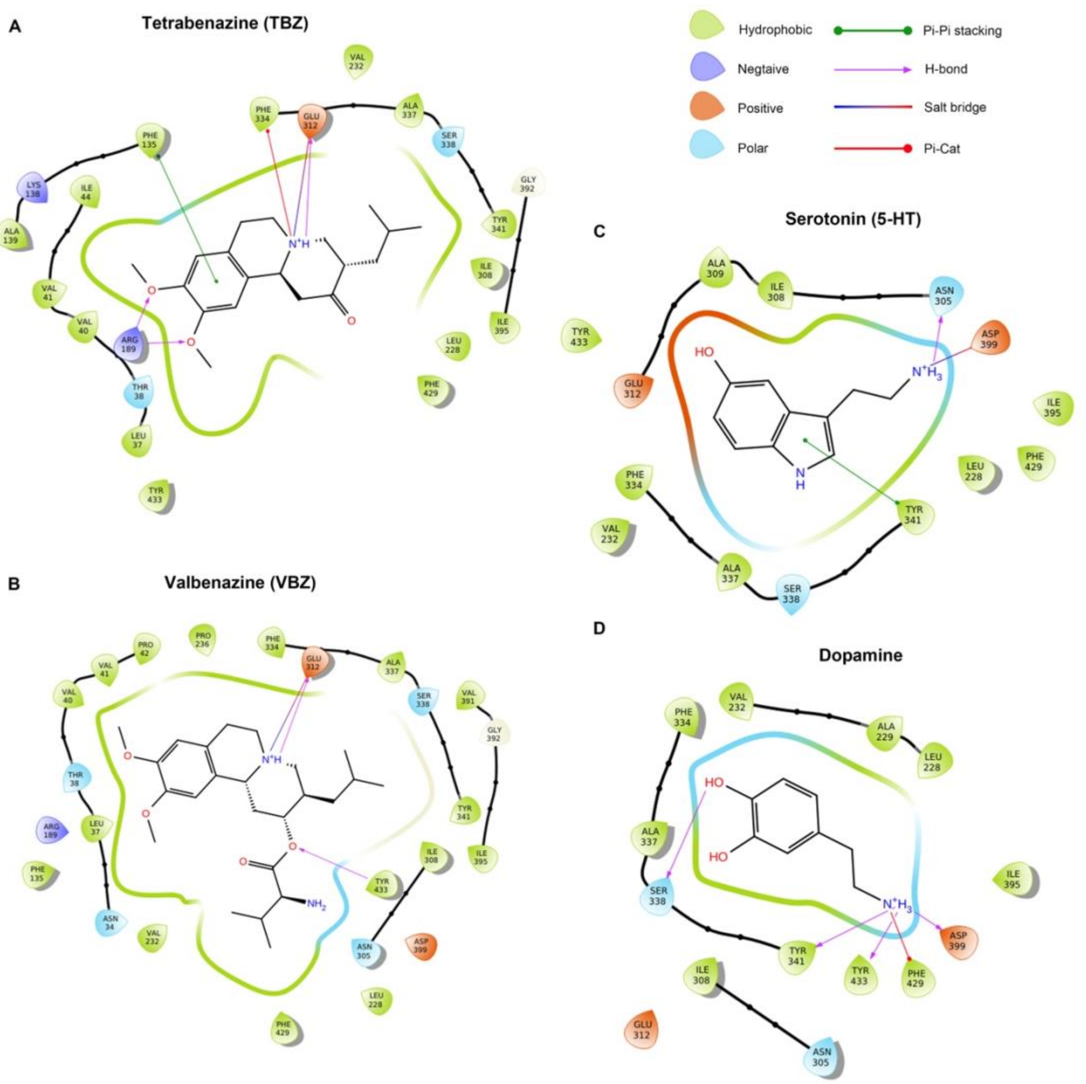

**Fig. S8 | Ligands-VMAT2 interaction diagrams.**

**A-D** Schematics of TBZ (A), VBZ (B), 5-HT (C), and DA (D) binding sites. Green, purple, orange, and blue drop shapes indicate hydrophobic, negative, positive, and polar residues, respectively. Ligand interactions are analyzed using Schrödinger Maestro.

**Table S1. Cryo-EM data collection, refinement and validation statistics**

| | VMAT2-TBZ (EMDB-38051) (PDB 8X4U) | VMAT2-VBZ (EMDB-38640) (PDB 8XT6) | VMAT2-5-HT (EMDB-38038) (PDB 8X3K) | VMAT2-DA (EMDB-38052) (PDB 8X4V) |
|---|---|---|---|---|
| **Data collection and processing** | | | | |
| Magnification | 130,000 | 130,000 | 130,000 | 105,000 |
| Voltage (kV) | 300 | 300 | 300 | 300 |
| Electron exposure ($e^-/Å^2$) | 48.23 | 49.42 | 49.97 | 49.5 |
| Defocus range (μm) | -0.9 to -1.6 | -0.9 to -1.6 | -0.9 to -1.6 | -1.3 to -1.8 |
| Pixel size (Å) | 0.932 | 0.932 | 0.932 | 0.832 |
| Symmetry imposed | C1 | C1 | C1 | C1 |
| Initial particle images (no.) | ~14.4 millions | ~18.8 millions | ~12.4 millions | ~6.9 millions |
| Final particle images (no.) | 143,834 | 146,307 | 214,255 | 52,161 |
| Map resolution (Å) | 3.46 | 3.48 | 2.98 | 3.63 |
| FSC threshold | 0.143 | 0.143 | 0.143 | 0.143 |
| **Refinement** | | | | |
| Initial model used (PDB code) | AlphaFold | 8X3K | AlphaFold | 8X3K |
| Map sharpening *B* factor ($Å^2$) | -188.0 | -180.0 | -145.9 | -144.9 |
| Model composition | | | | |
| Non-hydrogen atoms | 2783 | 2793 | 2797 | 2795 |
| Protein residues | 377 | 374 | 373 | 373 |
| Ligands | Tetrabenazine | Valbenazine | Serotonin | Dopamine |
| *B* factors ($Å^2$) | | | | |
| Protein | 124.88 | 89.66 | 74.94 | 123.92 |
| Ligand | 129.59 | 104.47 | 78.82 | 123.38 |
| R.m.s. deviations | | | | |
| Bond lengths (Å) | 0.005 | 0.007 | 0.008 | 0.007 |
| Bond angles (°) | 1.101 | 1.187 | 1.162 | 1.163 |
| Validation | | | | |
| MolProbity score | 1.98 | 1.98 | 1.78 | 1.81 |
| Clashscore | 12.21 | 13.55 | 8.39 | 10.84 |
| Poor rotamers (%) | 0.35 | 0.34 | 1.34 | 0.00 |
| Ramachandran plot | | | | |
| Favored (%) | 94.34 | 95.14 | 96.48 | 96.21 |
| Allowed (%) | 5.66 | 4.86 | 3.52 | 3.79 |
| Disallowed (%) | 0.00 | 0.00 | 0.00 | 0.00 |

**Table S2. Systems in MD simulations.**

| | |
|---|---|
| | — |
| | — |
| — | — |
| — | — |
| | — |
| | — |
| | — |
